# Interneuron-specific plasticity at parvalbumin and somatostatin inhibitory synapses onto CA1 pyramidal neurons shapes hippocampal output

**DOI:** 10.1101/774562

**Authors:** Matt Udakis, Victor Pedrosa, Sophie E.L. Chamberlain, Claudia Clopath, Jack R Mellor

## Abstract

The formation and maintenance of spatial representations within hippocampal cell assemblies is strongly dictated by patterns of inhibition from diverse interneuron populations. Although it is known that inhibitory synaptic strength is malleable, induction of long-term plasticity at distinct inhibitory synapses and its regulation of hippocampal network activity is not well understood. Here, we show that inhibitory synapses from parvalbumin and somatostatin expressing interneurons undergo long-term depression and potentiation respectively (PV-iLTD and SST-iLTP) during physiological activity patterns. Both forms of plasticity rely on T-type calcium channel activation to confer synapse specificity but otherwise employ distinct mechanisms. Since parvalbumin and somatostatin interneurons preferentially target perisomatic and distal dendritic regions respectively of CA1 pyramidal cells, PV-iLTD and SST-iLTP coordinate a reprioritisation of excitatory inputs from entorhinal cortex and CA3. Furthermore, circuit-level modelling reveals that PV-iLTD and SST-iLTP cooperate to stabilise place cells while facilitating representation of multiple unique environments within the hippocampal network.

## Introduction

GABAergic inhibitory interneurons form a diverse array of specialised cell types critical for the regulation of complex network functions within the brain. A defining feature of inhibitory interneurons is their precise axonal aborisations whereby inhibitory synapses target specific subdomains of pyramidal neurons and other inhibitory interneurons (Klausberger and Somogyi, 2008; Pelkey et al., 2017). Within the hippocampus and neocortex, parvalbumin (PV) and somatostatin (SST) expressing interneurons form two broad and occasionally overlapping subtypes of interneurons that preferentially target perisomatic and distal dendritic regions of pyramidal neurons respectively and are active on different phases of the theta cycle (Harris et al., 2018; Katona et al., 2014; Klausberger et al., 2003; Klausberger and Somogyi, 2008; Varga et al., 2012). This endows them with unique roles in sculpting pyramidal neuron responses to excitatory inputs (Milstein et al., 2015; Royer et al., 2012). Perisomatic inhibition by PV interneurons regulates pyramidal neuron spiking and network oscillations through feedforward and feedback inhibition (Geiger et al., 1997; Pouille and Scanziani, 2001; Sohal et al., 2009; Sun et al., 2014). In contrast, dendritic inhibition by SST interneurons regulates local synaptic and dendritic conductances, Ca^2+^ signal generation and excitatory synaptic plasticity principally through feedback inhibition (Chiu et al., 2013; Maccaferri, 2005; Schulz et al., 2018; Sun et al., 2014).

A defining feature of the hippocampus is the encoding of spatially relevant information via the formation of place cells (O’Keefe, 1976). Synaptic plasticity at glutamatergic synapses in the hippocampus accounts for the formation of location specific firing of individual place cells but it also plays a major role in the formation of place cell assemblies during exploration and offline replay of place cell activity (Bittner et al., 2017; Cohen et al., 2017; Isaac et al., 2009; Mishra et al., 2016; Sadowski et al., 2016). Interestingly, individual CA1 pyramidal neurons can encode distinct place fields in different environments (Colgin et al., 2008) presumably driven by ongoing excitatory synaptic plasticity. This feature of place cells and the persistent plasticity of their synaptic connections presents fundamental problems for hippocampal networks balancing flexibility versus stability of representations (Chaudhuri and Fiete, 2016). It is not clear how place cell assemblies in the hippocampus can encode multiple different locations in separate environments without interference.

Inhibitory interneurons play an integral role within the hippocampus controlling place cell activity (Del Pino et al., 2017; Royer et al., 2012; Sheffield et al., 2017; Trouche et al., 2016), where short-term changes in SST and PV interneuron activity differentially regulate the emergence and firing patterns of place cells (Royer et al., 2012; Sheffield et al., 2017) by controlling glutamatergic synaptic plasticity (Chiu et al., 2013; Schulz et al., 2018). But the consequences of long-term plasticity at inhibitory synapses on place cell activity has not been investigated.

Long-term inhibitory synaptic plasticity is a potentially important mechanism for learning within cortical networks (Chiu et al., 2019; Donato et al., 2013; Hellyer et al., 2016; Kullmann et al., 2012; Vogels et al., 2011) and GABAergic synapses in the hippocampus exhibit structural and functional plasticity (Muir et al., 2010; Nusser et al., 1998; Schuemann et al., 2013). Reductions in inhibitory strength via retrograde endocannabinoid signalling is well established (Alger and Pitler, 1995; Chevaleyre and Castillo, 2003; Lee et al., 2010) but multiple other mechanisms to regulate long-term inhibitory synaptic strength have also been proposed including GABA_B_ receptors and BDNF (Vickers et al., 2018), spike timing-dependent plasticity of chloride transporters (Woodin et al., 2003), retrograde nitric oxide signalling (Nugent et al., 2007) and NMDA receptors (Chiu et al., 2018). In the neocortex, synapses from PV and SST interneurons can undergo unique forms of plasticity (Chiu et al., 2018; Vickers et al., 2018), whilst in the hippocampus, recent evidence suggests interneuron subtype specific inhibitory synapses are regulated in distinct ways (Horn and Nicoll, 2018; Schulz et al., 2018). However, it is not clear whether long-term plasticity of inhibitory synapses is differentially engaged between interneuron subtypes during physiologically relevant activity and, furthermore, what the consequences of such plasticity would be for hippocampal network activity and place cell representations.

To address these questions, we utilised an optogenetic approach in hippocampal slices to selectively activate perisomatic and dendritically targeting inhibition onto CA1 pyramidal neurons by expression of channelrhodopsin in PV or SST interneurons. We found that synapses from PV and SST interneurons undergo interneuron-specific forms of inhibitory synaptic plasticity driven by the relative timing of inhibitory and excitatory neuronal spiking and employing distinct signalling mechanisms. We go on to show these forms of cell-specific long-term inhibitory plasticity have profound effects on the output of CA1 pyramidal neurons and use computational modelling to demonstrate that these plasticity rules can provide a mechanism by which hippocampal place fields can remain stable over time whilst flexibly encoding location in multiple environments.

## Results

### Divergent inhibitory plasticity at PV and SST synapses

To achieve subtype specific control of inhibitory interneurons, we selectively activated either PV or SST interneurons by expressing the light-activated cation channel channelrhodopsin-2 (ChR2) in a cre-dependent manner using mice that expressed cre recombinase under control of the promoter for either the parvalbumin gene (PV-cre) or somatostatin gene (SST-cre) crossed with mice expressing cre-dependent ChR2 (PV-ChR2 and SST-ChR2 mice; methods). Immunohistochemisty confirmed that ChR2 expression was highest in the Stratum Pyramidal (SP) and Stratum Oriens (SO) layers for PV-ChR2 mice with cell bodies principally located in SP (Figure 1A). Conversely, ChR2 expression was highest in the SO and Stratum Lacunosum Moleculare (SLM) layers for SST-ChR2 mice with cell bodies principally located in SO (Figure 1B). These expression profiles are consistent with the established roles of PV and SST interneurons providing perisomatic and dendritic inhibition respectively (Booker and Vida, 2018; Klausberger and Somogyi, 2008; Pelkey et al., 2017). To further confirm the spatially distinct inhibitory targets, we recorded interneuron subtype-specific inhibitory currents onto CA1 pyramidal neurons by activating ChR2 with 470nm blue light (Figure 1C). The rise and decay kinetics of the resulting light evoked PV IPSCs were significantly faster compared to SST derived IPSCs (Figure 1D,E) (Rise time: 3.8 ± 0.3 ms PV verses 6.1 ± 0.8 ms SST, n = 8, p < 0.05; Decay time: 16 ± 1.3 ms PV verses 28 ± 2.6 SST, n = 8, p < 0.01) supporting a more proximal location for PV synapses compared to SST synapses. Light evoked PV IPSC kinetics were almost identical to IPSC kinetics recorded from paired whole-cell recordings made from PV expressing fast-spiking basket cells to CA1 pyramidal cells (Figure S1A) and similarly, selective activation of Oriens Lacunosum Moleculare (OLM) cells using Chrna2-cre mice (Leao et al., 2012; Mikulovic et al., 2015) revealed IPSC kinetics indistinguishable from SST-ChR mice (Figure S1E). Furthermore, measurement of synaptic response amplitudes as light was targeted to different regions of the slice confirmed the immunohistochemical characterisation of ChR2 expression supporting the selective stimulation of perisomatic vs dendritic targeted inhibition (Figure S1B,C,F,G). Therefore, whilst we cannot exclude the activation of other interneuron subtypes expressing PV or SST, these data suggest the majority of our synaptic inputs most likely arise from activation of PV basket cells and SST OLM cells that selectively target synapses to perisomatic and distal dendritic regions of CA1 pyramidal cells respectively.

**Figure 1.**
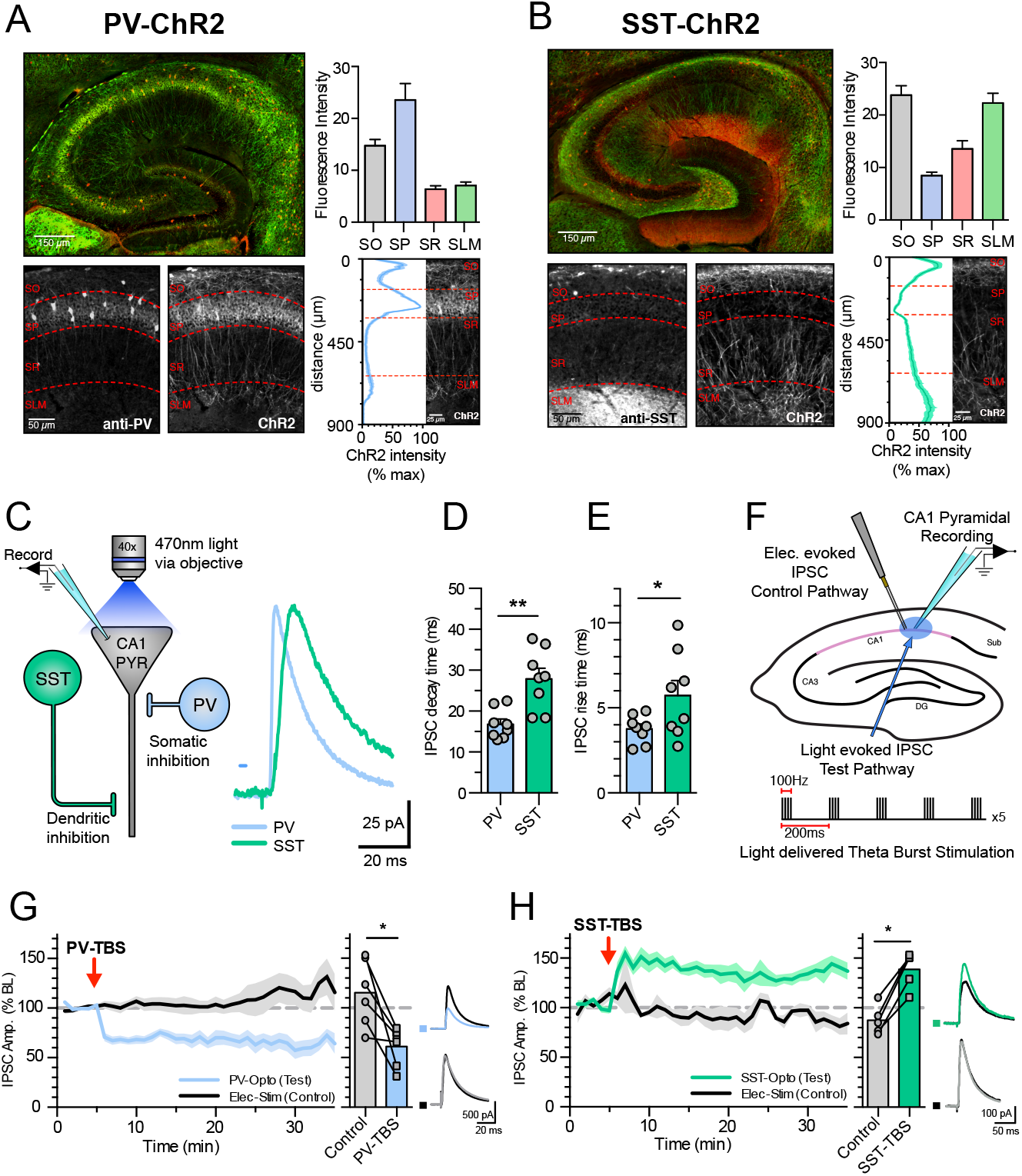
Somatically targeting PV and dendritically targeting SST inhibitory synapses undergo long-term synaptic plasticity. **(A)** Immunohistochemistry showing expression of PV (red) and ChR2 (green) in PV-ChR2 mice. Histogram displaying mean ChR2 fluorescence expression levels in different hippocampal layers: Stratum Oriens (SO), Stratum Pyramidal (SP) Stratum Radiatum (SR) and Stratum Lacunosum Moleculare (SLM) (right top), ChR2 expression as a function of distance across hippocampal layers (right bottom). **(B)** Same as A but for SST-ChR2 mice. **(C)** Schematic and example IPSC traces highlighting the somatic and distal targeting of PV and SST synapses. **(D)** Summary of IPSC decay times for PV and SST IPSCs. **(E)** Summary of IPSC rise times for PV and SST IPSCs. **(F)** Schematic displaying the recording set up for inhibitory plasticity experiments and the light induced theta burst (TBS) induction protocol. **(G)** TBS induced iLTD at PV synapses (left) average plasticity at control and test pathways (middle) and example traces before and after plasticity (right). **(H)** TBS induced iLTP at SST synapses (left) average plasticity at control and test pathways (middle) and example traces before and after plasticity (right). Data represent mean ± S.E.M statistical comparison via unpaired (D,E) and paired (G,H) t-tests where significance difference is indicated (*p < 0.05 and **p<0.01). See also Figures S1 and S2.

Having established two populations of inhibitory synapses we investigated whether PV or SST synapses undergo long-term inhibitory synaptic plasticity and if so, whether induction and expression is similar at each synapse. IPSCs were recorded from CA1 pyramidal neurons held at 0 mv with glutamatergic transmission pharmacologically blocked. Interneuron subtype-specific IPSCs mediated by GABA_A_ receptors were evoked by 5 ms light pulses and, importantly, an independent IPSC control pathway was evoked by electrical stimulation in the pyramidal layer (PV IPSCs) or Stratum Radiatum (SST IPSCs) (Figure 1F and Figure S1D,H). Both PV and SST interneurons are entrained to theta frequency rhythms in the hippocampus (Klausberger et al., 2003) so we first tested whether bursts of IPSCs delivered at theta frequency could induce long-term inhibitory synaptic plasticity. Theta burst stimulation (TBS) of the light evoked pathway led to a prolonged pathway-specific reduction of PV-IPSC amplitude indicating a synapse specific long-term depression of PV synapses (PV-iLTD, 115% ± 14% control verses 61% ± 8% test pathway, n = 6, p < 0.05) (Figure 1G). In contrast an identical light induced TBS led to a pronounced pathway-specific long-term potentiation of SST-IPSC amplitude (SST-iLTP, 87% ± 6% control verses 139% ± 8% test pathway, n = 6, p < 0.01) (Figure 1H). These findings indicate that high frequency inhibitory synaptic stimulation can induce inhibitory synaptic plasticity at PV and SST synapses, but the direction of plasticity is diametrically opposite for the two different synapses.

A common feature of plasticity at inhibitory synapses is the requirement for postsynaptic depolarisation in conjunction with synaptic stimulation, despite the synaptic stimulation itself causing hyperpolarisation (Alger and Pitler, 1995; Chevaleyre and Castillo, 2003; Chiu et al., 2018; Horn and Nicoll, 2018; Lee et al., 2010; Muir et al., 2010; Nusser et al., 1998; Schuemann et al., 2013; Schulz et al., 2018; Vickers et al., 2018; Woodin et al., 2003). To test the requirement for depolarisation in PV-iLTD and SST-iLTP we delivered TBS whilst voltage clamping neurons at −60mV during the TBS protocol. Under these conditions neither PV-iLTD nor SST-iLTP were induced (Figure S2A,B; PV: 84% ± 8% control verses 91% ± 4% test pathway, n = 5, p > 0.05; SST: 96% ± 6% control verses 106% ± 4% test pathway, n = 5, p > 0.05), indicating that both forms of inhibitory plasticity require coincident pyramidal neuron depolarisation and inhibitory input.

### PV and SST synapses undergo spike timing-dependent inhibitory plasticity

During exploratory behaviour, neurons in the hippocampus are entrained to the theta rhythm (Buzsaki, 2002) with defined populations of neurons, including different subpopulations of interneurons, firing action potentials at specific phases of the theta cycle (Klausberger et al., 2003; Klausberger and Somogyi, 2008; Varga et al., 2012). Having established a requirement for coincident synaptic activity and postsynaptic depolarisation for the induction of PV-iLTD and SST-iLTP we next sought to determine whether this was a Hebbian form of plasticity that could be induced by coincident pre- and post-synaptic action potentials and if so, what the precise spike timing requirements might be with respect to the preferred interneuron and pyramidal spiking phases of the theta cycle. The major subclasses of PV interneurons, basket cells and axo-axonic cells, fire on the descending phase of theta cycle roughly 60ms before pyramidal neurons, whilst both bistratified and OLM SST interneurons fire near coincident with pyramidal neurons at the trough of the cycle (Figure 2A) (Katona et al., 2014; Klausberger et al., 2003; Klausberger and Somogyi, 2008; Varga et al., 2012). We therefore tested the induction of inhibitory spike timing-dependent plasticity (iSTDP) using spike timings of −60 ms, 0 ms and +60 ms to replicate spike patterns during exploratory behaviour and span the full width of a theta cycle (Figure 2A).

**Figure 2.**
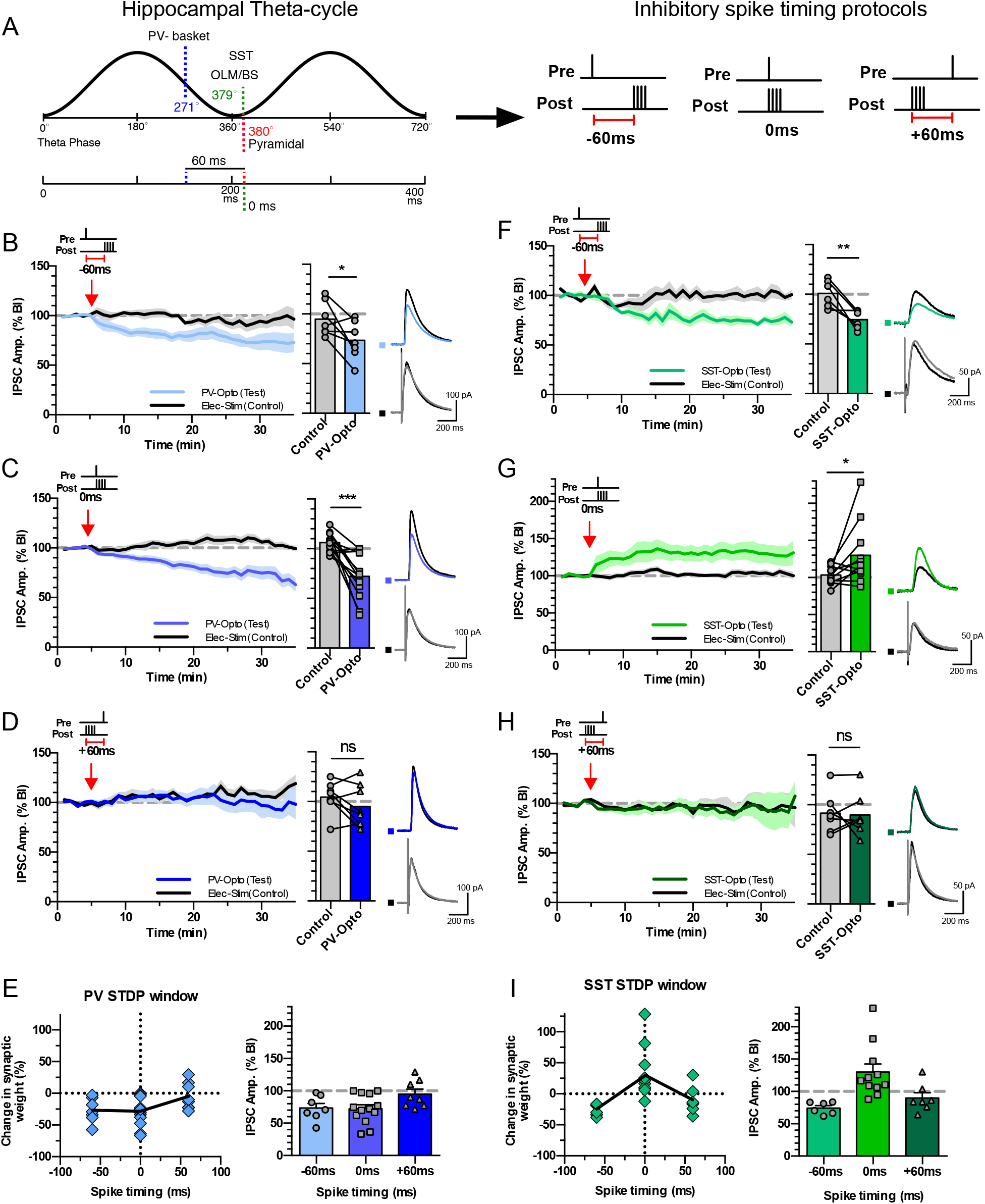
PV and SST inhibitory synapses undergo spike timing-dependent plasticity. **(A)** Schematic highlighting the relative spike timing of PV and SST expressing interneurons during theta oscillations in relation to pyramidal neuron spiking (left). The three pairing protocols used for iSTDP experiments representing presynaptic stimulation and postsynaptic action potentials −60 ms pre before post, 0 ms pre and post together and +60 ms post before pre (right). **(B)** −60 ms pre before post pairing induced iLTD at PV synapses (left) average plasticity at control and test pathways (middle) and example traces for before and after plasticity (right). **(C)** 0 ms pre and post pairing induced iLTD at PV synapses (left) average plasticity at control and test pathways (middle) and example traces for before and after plasticity (right). **(D)** +60 ms post before pre pairing failed to induce plasticity at PV synapses (left) average plasticity at control and test pathways (middle) and example traces for before and after plasticity (right). **(E)** Summary of the inhibitory STDP window at PV synapses. **(F,G,H,I)** Same as B,C,D,E but for SST synapses. Data represent mean ± S.E.M statistical comparison via paired t-tests where significance difference is indicated (*p < 0.05 and ***p<0.001). Scale bars B-D: 200ms, 100pA F-H 200ms, 50pA.

To test iSTDP, CA1 pyramidal neurons were voltage clamped at −50mV in the presence of AMPA and NMDA receptor blockers and light and electrically evoked IPSCs recorded as test and control synaptic pathways respectively. During the iSTDP protocol, recordings were switched to current clamp with the membrane potential maintained at −50 mV and single light evoked IPSCs were paired with a burst of 4 action potentials repeated 100 times at theta frequency (5 Hz). Stimulation of PV synapses with the theta-relevant −60 ms spike timing protocol resulted in pathway specific PV-iLTD (Figure 2B, 95% ± 7% control verses 73% ± 7% test pathway, n = 7, p < 0.05). PV-iLTD was also observed when PV synapses were paired at 0 ms (Figure 2C, 106% ± 3% control verses 72% ± 6% test pathway, n = 13, p < 0.001) but not at +60 ms (Figure 2D, 105% ± 6% control verses 95% ± 8% test pathway, n = 8, p > 0.05) resulting in a pan-theta cycle iSTDP relationship incorporating iLTD but no iLTP (Figure 2E). Surprisingly, SST synapses also displayed iLTD at −60 ms spike timings (Figure 2F, 101% ± 7% control verses 70% ± 5% test pathway, n = 6, p < 0.01) but contrastingly underwent iLTP when paired at 0 ms (Figure 2G, 104% ± 4% control verses 130% ± 13% test pathway, n = 11, p < 0.05). At pairings of +60 ms SST synapses also exhibited no plasticity similar to PV synapses (Figure 2H, 92% ± 8% control verses 90% ± 8% test pathway, n = 7, p > 0.05). Therefore, SST synapses can undergo both iLTD and iLTP depending on the precise spike timing of pre- and post-synaptic action potentials in contrast to PV synapses that only undergo iLTD. These results demonstrate that spike timings observed during theta rhythm entrainment lead to distinct rules for iSTDP at PV and SST synapses. Interestingly, we observed at both PV and SST synapses that pairing inhibitory inputs 60 ms after a burst of action potentials was insufficient to induce inhibitory plasticity, highlighting the importance of spike timing and the need for inhibitory synaptic input prior to pyramidal neuron activity.

### PV-iLTD requires postsynaptic activation of T-type Ca^2+^ channels and calcineurin

We next investigated the molecular mechanisms of spike timing-dependent PV-iLTD. First, we found that presynaptic input or postsynaptic spikes alone were insufficient to induce iLTD at PV synapses. (Presynaptic input only: 100% ± 12% control verses 94% ± 6% test pathway, n = 6, p > 0.05; Postsynaptic spikes only: 103% ± 10% control verses 94% ± 5% test pathway, n = 6, p > 0.05) (Figure 3A,B). Consistent with PV-iSTDP and TBS induced PV-iLTD, these results show that coincident activity between PV interneurons and pyramidal neurons is required for PV-iLTD. Many forms of synaptic plasticity also depend on elevations in postsynaptic Ca^2+^ and we tested if this was the case for PV-iLTD by including the Ca^2+^ chelator BAPTA in the intracellular recording solution. BAPTA prevented PV-iLTD demonstrating a dependence on postsynaptic Ca^2+^ (100% ± 10% control verses 95% ± 8% test pathway, n = 6, p > 0.05) (Figure 3C).

**Figure 3.**
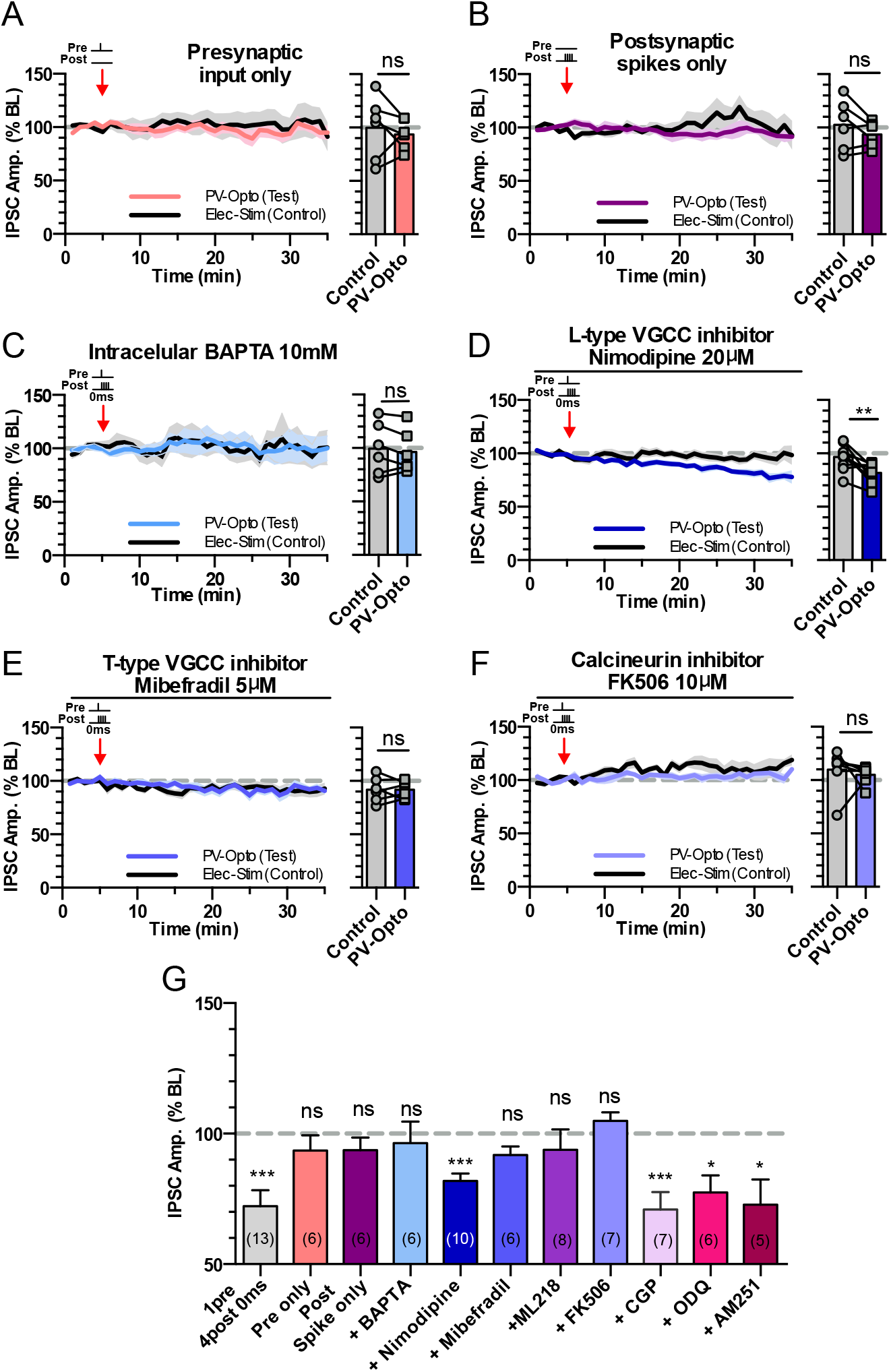
PV-iLTD requires activation of T-type VGCCs and calcineurin. **(A)** Presynaptic stimulation of PV inputs alone induced no plasticity. **(B)** Post synaptic spikes alone failed to induce plasticity at PV synapses. **(C)** inclusion of BAPTA in the internal recording solution occludes PV iLTD upon 0 ms pre and post pairing. **(D)** L-type calcium channel antagonist Nimodopine doesn’t block PV iLTD upon 0 ms pre and post pairing. **(E)** T-type calcium channel antagonist Mibefradil occludes PV iLTD upon 0 ms pre and post pairing. **(F)** Calcineurin inhibitor, FK506 occludes PV iLTD upon 0 ms pre and post pairing. In panels A-F, average plasticity in control and test pathways is shown on the right. **(G)** Summary histogram displaying the level of plasticity under each experimental condition. Data represent mean ± S.E.M statistical comparison via paired t-tests (A-F) and one sample t-tests (G). Significant difference is indicated (*p < 0.05, **p<0.01 and ***p<0.001). See also Figure S3.

Important sources of postsynaptic Ca^2+^ for the induction of excitatory and inhibitory synaptic plasticity are NMDA receptors and voltage-gated Ca^2+^ channels (VGCCs) (Chiu et al., 2018; Griffith et al., 2016; Magee and Johnston, 1997; Tigaret et al., 2016). Since NMDA receptors are blocked in our experiments, we investigated the role of VGCCs in PV-iLTD. L-type VGCCs are the most prominent postsynaptic VGCCs, however, the L-type VGCC inhibitor nimodipine (20 µM) failed to block PV-iLTD (97% ± 4% control verses 82% ± 3% test pathway, n = 10, p < 0.01 (Figure 3D). We next tested the role of T-type VGCCs using the inhibitor mibefradil (5 µM) which blocked PV-iLTD (92% ± 5% control verses 92% ± 3% test pathway, n = 6, p > 0.05 (Figure 3E) and this was confirmed with the use of another T-type VGCC inhibitor ML218 (3 µM) (Xiang et al., 2011) (99.4% ±5.3% control verses 93.8 ± 7.2% test pathway, n = 8, p > 0.05) (Figure S3E). Interestingly, T-type VGCCs have a low voltage threshold for activation and predominantly reside in an inactivated state at resting membrane potentials (Randall and Tsien, 1997). They therefore require hyperpolarisation to relieve voltage inactivation (de-inactivation), which corresponds precisely with the requirement for inhibitory synaptic input prior to postsynaptic depolarisation resulting in synapse specificity of PV-iLTD. These findings highlight a mechanism by which inhibitory synapses can provide a synapse-specific source of Ca^2+^ to induce inhibitory plasticity.

The downstream effects of Ca^2+^ can lead to release of retrograde signalling molecules which regulate presynaptic release of GABA (Chevaleyre and Castillo, 2003; Nugent et al., 2007) or it can signal postsynaptically to reduce GABA_A_ receptor function (Muir et al., 2010). We therefore tested whether previously described retrograde signalling molecules nitrous oxide (Nugent et al., 2007) and endocannabinoids (Chevaleyre and Castillo, 2003) are involved in PV-iLTD. The nitrous oxide pathway antagonist ODQ (5 µM) and the CB1 receptor antagonist AM251 (1 µM) both failed to prevent PV-iLTD (Figure S3C,D and Figure 3G, ODQ: 100% ± 8% control verses 78% ± 7% test pathway, n = 6, p < 0.05; AM251: 98% ± 8% control verses 73% ± 10% test pathway, n = 5, p < 0.05). An additional candidate mechanism could be the activation of the G-protein coupled GABA_B_ receptor since this has been shown to mediate a form of iLTD (Vickers et al., 2018), however the GABA_B_ receptor antagonist CGP 55845 (1 µM) also failed to prevent PV-iLTD (Figure S3B and Figure 3G, 101% ± 10% control verses 71% ± 7% test pathway, n = 7, p < 0.05). One postsynaptic signalling pathway that has been implicated in iLTD is activation of the phosphatase calcineurin (Luscher et al., 2011; Muir et al., 2010; Wang et al., 2003). Application of the calcineurin inhibitor FK506 (10 µM) prevented PV-iLTD (110% ± 7% control verses 105% ± 3% test pathway, n = 7, p > 0.05) (Figure 3F) indicating a postsynaptic target for Ca^2+^ signalling. In summary, PV-iLTD requires coincident pre- and post-synaptic activity, opening T-type VGCC to provide a postsynaptic Ca^2+^ signal that leads to activation of calcineurin to induce LTD at PV inhibitory synapses (Figure 3G, 5A).

### SST-iLTP requires postsynaptic activation of L- and T-type Ca^2+^ channels and CAMKII

The molecular mechanisms of spike timing-dependent SST-iLTP were next investigated. Similar to PV-iLTD, and consistent with SST-iSTDP and TBS induced SST-iLTP, we found that SST-iLTP requires coincident pre- and post-synaptic activation as neither SST inputs nor postsynaptic action potentials alone were able to induce SST-iLTP (Presynaptic input only: 107% ± 6% control verses 91% ± 11% test pathway, n = 6, p > 0.05; Postsynaptic spikes only: 94% ± 5% control verses 100% ± 10% test pathway, n = 7, p > 0.05) (Figure 4A,B). The inclusion of the Ca^2+^ chelator BAPTA also prevented the induction of iLTP at SST synapses (97.44% ± 10% control verses 92% ± 6% test pathway, n = 6, p > 0.05) (Figure 4C), indicating SST-iLTP requires postsynaptic Ca^2+^. Again, we assessed if L- and/or T-type VGCCs could provide the source of postsynaptic Ca^2+^ required for SST-iLTP. Interestingly, L-Type VGCC antagonist nimodipine and T-type VGCC antagonists mibefradil and ML218 both blocked SST-iLTP (nimodipine: 93% ± 7% control verses 101% ± 7% test pathway, n = 5, p > 0.05; mibefradil: 99% ± 9% control verses 101% ± 10% test pathway, n = 7, p > 0.05; ML218: 113% ± 8% control verses 100% ± 6% test pathway, n = 7, p > 0.05) (Figure 4D,E,H, Figure S4B) showing SST-iLTP requires activation of L-type and T-type VGCCs with the latter providing a synapse specific source of Ca^2+^ similar to PV-iLTD.

**Figure 4.**
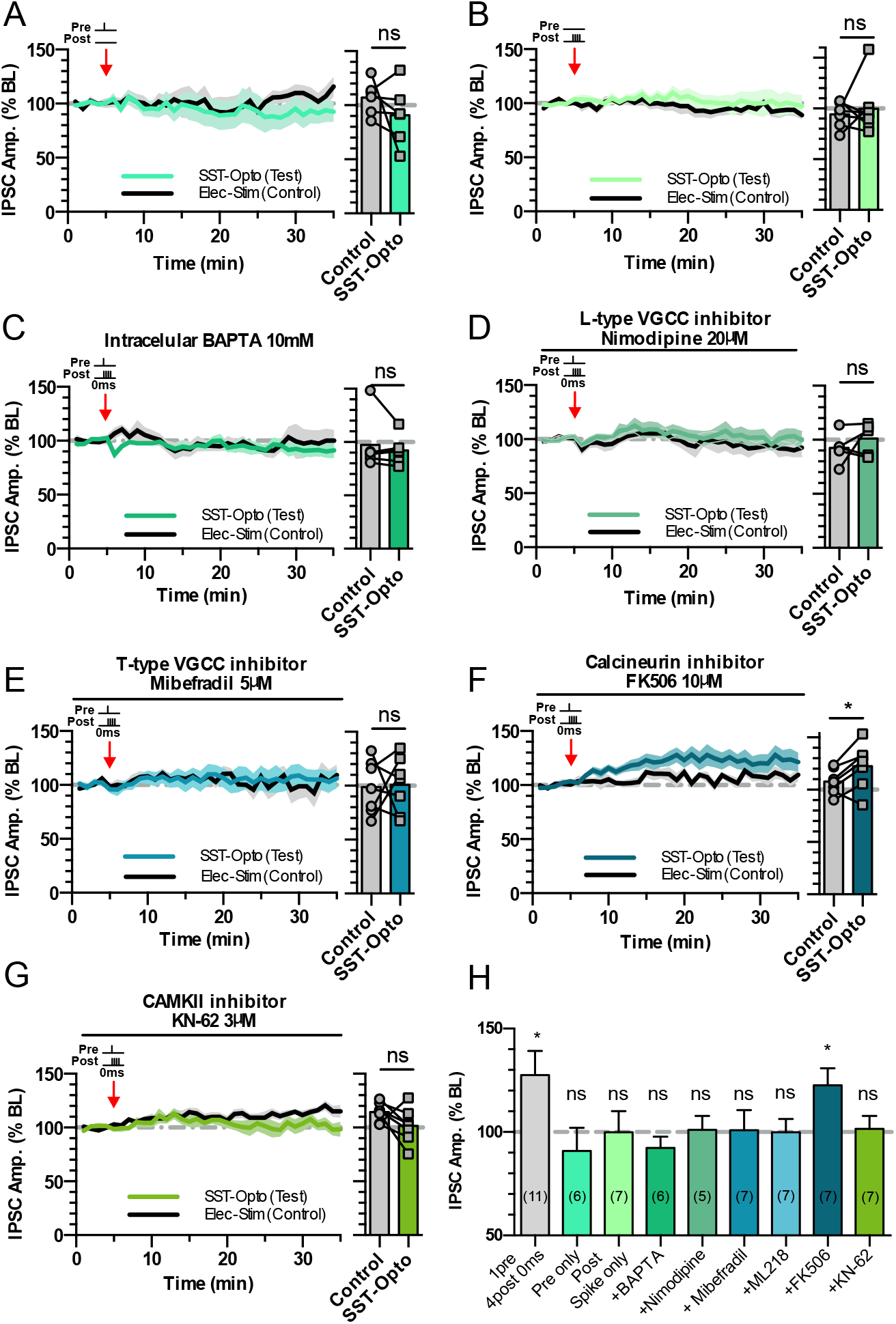
SST-iLTP requires activation of L-type and T-type VGCCs and CAMKII. **(A)** Presynaptic stimulation of SST inputs alone failed to induce plasticity. **(B)** Post synaptic spikes alone failed to induce plasticity at SST synapses. **(C)** Inclusion of BAPTA in the internal recording solution occludes SST iLTP upon 0 ms pre and post pairing. **(D)** L-type calcium channel antagonist Nimodopine occludes SST iLTP upon 0 ms pre and post pairing. **(E)** T-type calcium channel antagonist Mibefradil occludes SST iLTP upon 0 ms pre and post pairing. **(F)** Calcineurin inhibitor, FK506 fails to block SST iLTP upon 0 ms pre and post pairing. **(G)** CAMKII inhibitor KN-62 occludes SST iLTP upon 0 ms pre and post pairing. In panels A-G, average plasticity in control and test pathways is shown on the right. **(H)** Summary histogram displaying the level of plasticity under each experimental condition. Data represent mean ± S.E.M statistical comparison via paired t-tests (A-G) and one sample t-tests (H). Significant difference is indicated (*p < 0.05). See also Figure S4.

Since calcineurin is required for PV-iLTD, we next tested the possible involvement of calcineurin in SST-iLTP. However, inhibition of calcineurin failed to block SST-iLTP (108% ± 5% control verses 123% ± 8% test pathway, n = 7, p < 0.05) (Figure 4F,H). As SST-iLTP requires both L-type and T-type VGCCs we hypothesised that SST-iLTP might be mediated by a molecular pathway engaged by high levels of postsynaptic Ca^2+^. CAMKII is one such candidate and has been shown to mediate potentiation of inhibitory synapses including SST synapses within the cortex (Chiu et al., 2018) and other inhibitory synapses within the hippocampus (Marsden et al., 2007; Petrini et al., 2014) resulting in postsynaptic changes in GABA_A_ receptors. We therefore tested the involvement of CAMKII activation on hippocampal SST-iLTP and found that the CAMKII inhibitor KN-62 (3 µM) prevented SST-iLTP (115% ± 3% control verses 102% ± 6% test pathway, n = 7, p > 0.05) (Figure 4G,H), consistent with its role in mediating iLTP. In summary, SST-iLTP requires the coincident activation of SST synapses and pyramidal neurons, which activates L-type and T-type VGCCs providing a Ca^2+^ source able to activate CAMKII to induce SST-iLTP (Figure 5B).

**Figure 5.**
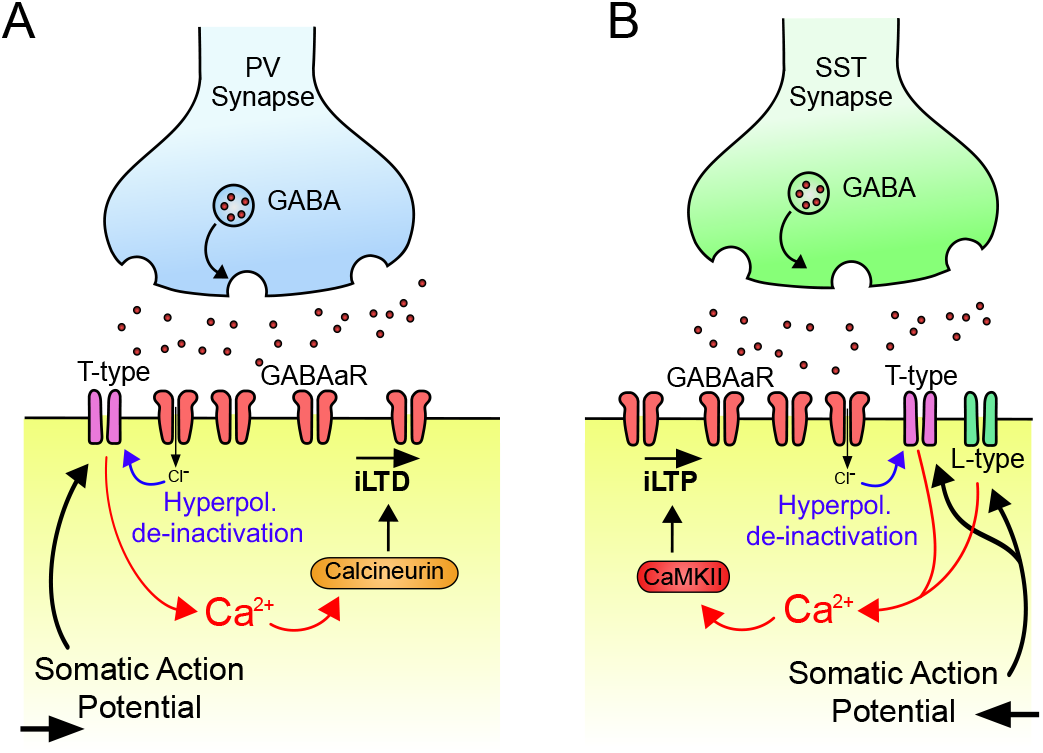
PV-iLTD and SST-iLTP mechanisms. **(A)** Illustration of PV-iLTD mechanism. Hyperpolarisation by GABAA receptor currents relieves T-type VGCCs from voltage dependent inactivation (de-inactivation). Back-propagating action potentials then activate T-type VGCCs providing an inhibitory synapse specific source of Ca2+ to activate calcineurin resulting in LTD at PV synapses. **(B)** Illustration of SST-iLTP mechanism. Similar to PV-iLTD but requiring activation of T-type and L-type VGCCs that activates CAMKII resulting in LTP at SST synapses.

### SST and PV plasticity shape pyramidal neuron responses to excitatory input pathways

To understand the potential implications of PV-iLTD and SST-iLTP on network integration of inputs to a pyramidal neuron, we implemented a multi-compartment model of a CA1 pyramidal neuron in the presence of proximal (PV) and distal (SST) inhibition (Figure 6A). The simulated CA1 pyramidal neuron receives distal excitatory input from entorhinal inputs via the temporoammonic (TA) pathway and proximal excitatory input from CA3 inputs via the Schaffer collateral (SC) pathway. We also implemented rate-based inhibitory plasticity rules derived from our experiments under the most physiologically-relevant conditions (SST-iLTP and PV-iLTD). Inhibitory synaptic weights onto pyramidal cells were therefore updated following a Hebbian plasticity rule in which coincident pre- and postsynaptic activity leads to iLTD for PV synapses and iLTP for SST synapses. We then simulated activity within the network before and after the induction of inhibitory plasticity and compared the correlation between Schaffer collateral or temporoammonic inputs and pyramidal cell activity at these two stages. As expected, the correlation between Schaffer collateral inputs and CA1 pyramidal cell activity increased and the correlation between temporoammonic inputs and CA1 pyramidal cell activity decreased following the induction of interneuron plasticity (Figure 6B). Therefore, if we assume that SST-iLTP occurs primarily at distal dendritic locations, interneuron-specific plasticity is a potential mechanism to change CA1 network state from being driven by both temporoammonic and Schaffer collateral inputs to being primarily driven by Schaffer collateral inputs.

**Figure 6.**
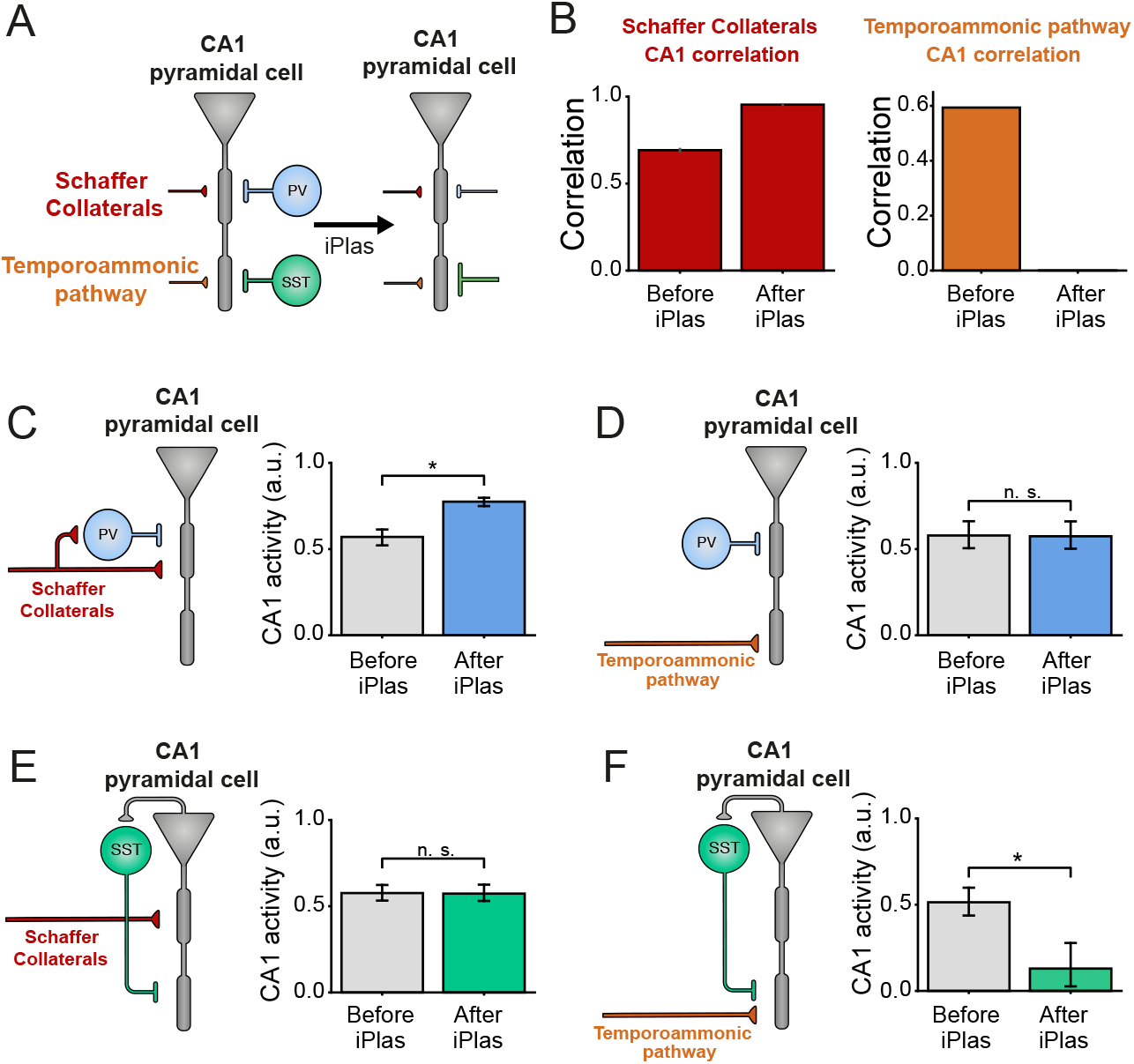
PV and SST plasticity differentially regulate Schaffer collateral and temporoammonic excitation of CA1 pyramidal neurons. **(A)** Diagram of a simulated, rate-based CA1 pyramidal cell before and after the induction of inhibitory plasticity (iPlas). A single two-compartment neuron receives inputs from four sources: distally-targeting temporoammonic, proximally-targeting Schaffer collaterals, distally-targeting inhibition from SST interneurons, and proximally-targeting inhibition from PV interneurons. iPlas leads to PV-iLTD and SST-iLTP. **(B)** Correlation between Schaffer collateral activity and CA1 somatic activity (left) and temporoammonic activity and CA1 somatic activity (right) before and after iPlas at PV and SST synapses (PV-iLTD and SST-iLTP). **(C-F)** CA1 somatic activity before and after iPlas (either PV-iLTD or SST-iLTP) under the individual stimulation of either Schaffer collaterals or temporoammonic inputs. **(C)** Schaffer collateral-driven CA1 somatic activity is enhanced upon PV-iLTD. **(D)** CA1 somatic activity driven by temporoammonic input after PV-iLTD, is unchanged. **(E)** CA1 somatic activity driven via Schaffer collateral input after SST-iLTD is unchanged. **(F)** SST-iLTP leads to a reduction in CA1 somatic activity in response to temporoammonic input. See also Figure S5. Data represent mean ± S.E.M statistical comparison via unpaired t-tests. Significant difference is indicated (*p < 0.05).

We next incorporated functionally-relevant feedforward and feedback connectivity for PV and SST interneurons within the CA1 network. PV interneurons receive strong feedforward innervation from the Schaffer collateral pathway but relatively limited input from the temporoammonic pathway and some feedback input from CA1 pyramidal neurons (Booker and Vida, 2018; Ganter et al., 2004; Milstein et al., 2015; Sun et al., 2014). In contrast, distally targeting SST interneurons receive almost no feedforward input and are driven by feedback input from CA1 pyramidal neurons (Blasco-Ibanez and Freund, 1995; Booker and Vida, 2018; Lacaille et al., 1987; Sun et al., 2014). There is also evidence that bistratified interneurons that target inhibition to proximal dendrites in Stratum Radiatum and express both PV and SST can be feedforward in the Schaffer collateral pathway and also feedback within CA1 (Booker and Vida, 2018; Sun et al., 2014).

Using these various functional connectivity arrangements, we first investigated the consequences of PV-iLTD on CA1 pyramidal cell output. For Schaffer collateral inputs with feedforward PV interneurons, PV-iLTD led to an increase in CA1 pyramidal cell activity due to a reduction in feedforward inhibition (Figure 6C). The same result was achieved if we included feedback inhibition from PV interneurons (Figure S5A). When we considered the temporoammonic pathway without feedforward PV interneurons, PV-iLTD did not change pyramidal cell output (Figure 6D). However, if we incorporated PV interneurons as feedforward and feedback inhibition, PV-iLTD led to an increase in pyramidal cell activity in response to temporoammonic stimulation (Figure S5B). Therefore, our model predicts that CA1 pyramidal cell activity in response to Schaffer collateral stimulation should increase following PV-iLTD whereas it is likely to remain unchanged in response to stimulation of the temporoammonic pathway unless PV interneurons participating in feedforward inhibition of the temporoammonic pathway or feedback inhibition are significantly activated and undergo PV-iLTD.

We next investigated the implications of SST-iLTP on CA1 pyramidal cell output. Pyramidal cell activity in response to Schaffer collateral stimulation was not affected by SST-iLTP if SST synapses are located at the pyramidal cell’s distal dendritic compartment (Figure 6E) and this was true regardless of whether SST interneurons were activated in feedforward or feedback fashion (Figure S5C). Contrary to Schaffer collateral stimulation, temporoammonic-induced CA1 pyramidal cell activity was reduced after SST-iLTP (Figure 6F) and this effect was stronger if SST interneurons were considered to be feedforward as well as feedback (Figure S5D). In summary, our model predicts that SST-iLTP does not affect Schaffer collateral-induced CA1 pyramidal cell activity whereas SST-iLTP decreases activity induced by temporoammonic stimulation.

If we assume the functional connectivity shown in Figures 6C-F, our model simulations therefore predict that PV-iLTD will increase CA1 pyramidal neuron responses to Schaffer collateral but not temporoammonic inputs and that SST-iLTP will decrease CA1 pyramidal neuron responses to temporoammonic but not Schaffer collateral inputs.

To test these predictions, we experimentally investigated the impact of PV-iLTD and SST-iLTP on the probability of spike generation in CA1 pyramidal neurons. By stimulating either the Schaffer collateral or temporoammonic pathways, action potential probability was recorded in response to 10 consecutive EPSPs where the stimulation intensity was set such that the baseline action potential probability for each EPSP was ∼50%. Upon Schaffer collateral stimulation, PV-iLTD (0 ms timing) led to an increase in the spike probability (0.4 ± 0.07 baseline verses 0.78 ± 0.05 post plas, n = 6, p < 0.05) (Figure 7A) that mirrored the timecourse of PV-iLTD, but for temporoammonic pathway stimulation spike probability was unaltered (0.44 ± 0.04 baseline verses 0.47 ± 0.03 post plas, n = 6, p > 0.05) (Figure 7B). Taken together these results confirm the predictions from the model and suggest that in our experiments the majority of PV interneurons recruited by Schaffer collateral stimulation are feedforward and undergo PV-iLTD whereas few PV interneurons are recruited by temporoammonic stimulation or via feedback excitation from CA1 pyramidal neurons. We next conducted experiments to examine the impact of SST-iLTP on spike generation. We found that the increase in SST inhibition with SST-iLTP had little effect on action potential generation from Schaffer collateral stimulation (0.53 ± 0.04 baseline verses 0.5 ± 0.08 post plas, n = 8, p > 0.05) (Figure 7C) but led to a robust reduction in spike generation from the temporoammonic pathway (0.52 ± 0.03 baseline verses 0.27 ± 0.09 post plas, n = 6, p < 0.05 (Figure 7D). Again, these results confirm the predictions from the model and suggest that in our experiments SST interneurons are primarily feedback and target distal dendritic regions of pyramidal neurons.

**Figure 7.**
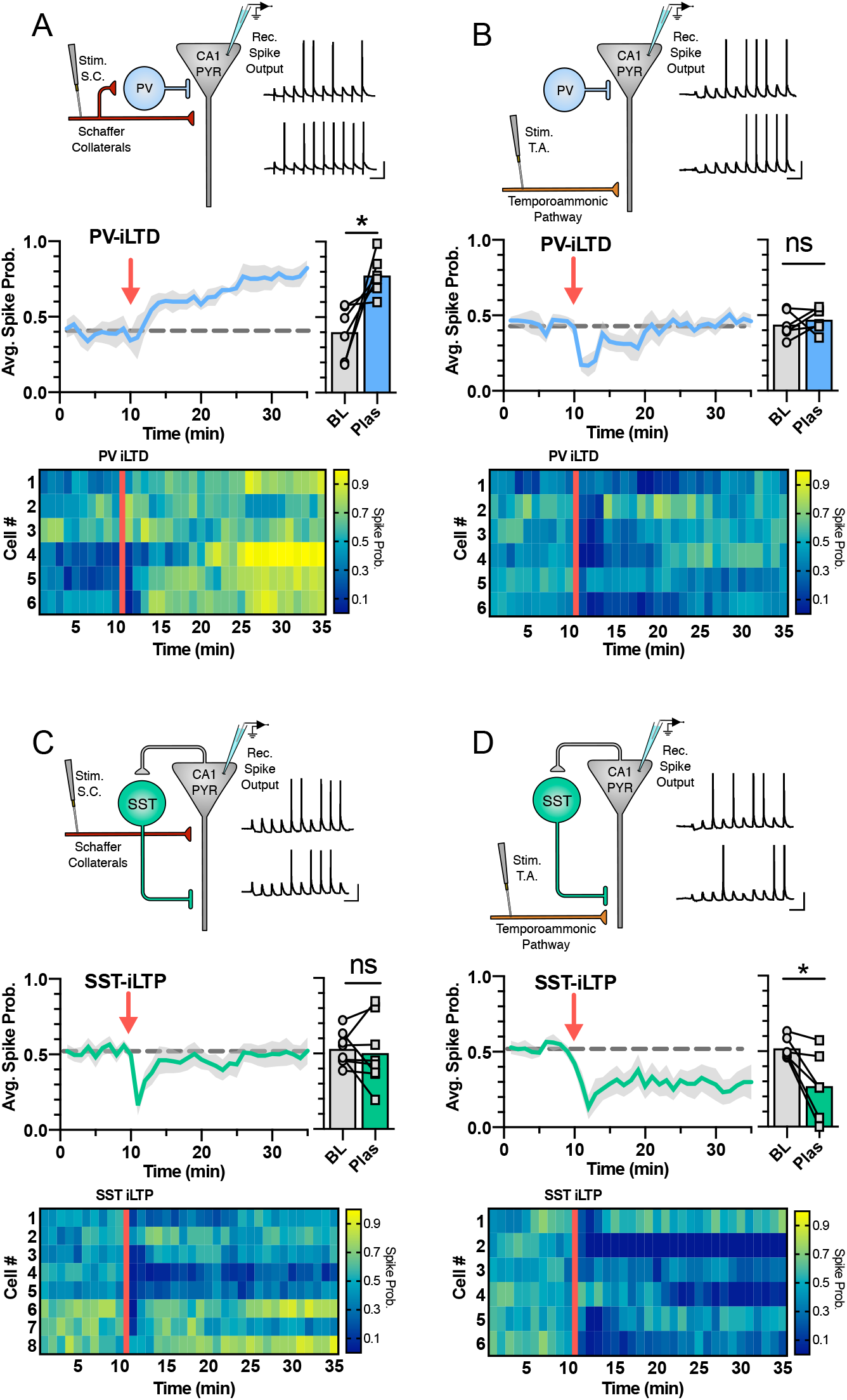
CA1 output driven by Schaffer collateral or temporoammonic inputs is differentially regulated by PV-iLTD and SST-iLTP. **(A)** Diagram showing the experimental design where electrically stimulated Schaffer collaterals evoke action potentials in CA1 pyramidal neurons with example current-clamp traces before and after induction of PV iLTD (0 ms pre and post pairing) (top). Average spike probability during baseline (BL) and 20-25 minutes after induction of PV iLTD (Plas) (middle). Minute average data for each cell showing the spike probability during baseline and after PV iLTD. **(B)** Same as A but for the stimulation of the temporoammonic pathway. **(C)** Diagram showing electrical stimulation of Schaffer collaterals with example traces before and after induction of SST iLTP (0 ms pre and post pairing) (top). Average spike probability during baseline (BL) and after SST iLTP (Plas) (middle). Minute average data for each cell showing the spike probability during baseline and after SST iLTP. **(D)** same as C but for stimulation of the temporoammonic pathway. Data represent mean ± S.E.M statistical comparison via paired t-tests between baseline and post plasticity where significance difference is indicated (*p < 0.05). Scale bars: 100ms, 20mV.

Taken together, these results demonstrate that long-term inhibitory plasticity changes the responses of CA1 pyramidal neurons prioritising inputs from the Schaffer collateral pathway over those from the temporoammonic pathway. Increased distal dendritic inhibition driven by SST-iLTP will also inhibit the induction of excitatory synaptic plasticity (Leao et al., 2012; Williams and Holtmaat, 2019) with important functional implications for the formation and stability of place cells.

### Inhibitory plasticity at PV and SST synapses provides a mechanism for place cell stability across multiple environments

We next explored the implications of long-term inhibitory plasticity on place cell physiology within hippocampal networks. The long-term nature of inhibitory plasticity suggests that its impact on place cell physiology will be evident as an animal traverses different environments. Key features of place cells are: (i) that in multiple different environments each place cell may represent distinct locations or switch to be silent, and (ii) that within any single environment place cell representations are broadly stable upon repeated exposures to that environment (Colgin et al., 2008; Ziv et al., 2013). However, these two features are somewhat contradictory since they require place cells to respond to different inputs without interference (Chaudhuri and Fiete, 2016).

To investigate the functional implications of interneuron subtype-dependent long-term plasticity for place cell physiology in multiple different environments, we simulated a CA1 network receiving place-tuned input while an animal explored first an annular track (environment A) and then a different track (environment B) before finally returning to the original familiar environment A’ (Figure 8A). In our simulations, CA1 pyramidal neurons receive inputs from Schaffer collaterals, temporoammonic pathway, SST interneurons and PV interneurons (Figure 8B). Schaffer collateral inputs are spatially tuned while the other inputs are considered spatially uniform for simplicity. Schaffer collateral inputs are plastic and follow a Hebbian-type plasticity rule which depends on coincident pre- and post-synaptic activation. Following recent evidence that place fields are formed by synaptic plasticity at Schaffer collateral synapses following closely-timed temporoammonic and Schaffer collateral inputs (Bittner et al., 2015; Bittner et al., 2017), we implemented the postsynaptic term of the Hebbian plasticity rule to be the product of the activities of the distal and proximal compartments of our two-compartment neuron model (see Methods). SST and PV inhibitory synapses onto pyramidal cells also follow a rate-based Hebbian-type plasticity rule inspired by the physiologically-relevant scenarios from our experimental data (as for Figure 6). Coincident pre- and post-synaptic activity leads to iLTP in the case of SST synapses, whereas pre- and post-synaptic coactivation leads to iLTD in the case of PV synapses.

**Figure 8.**
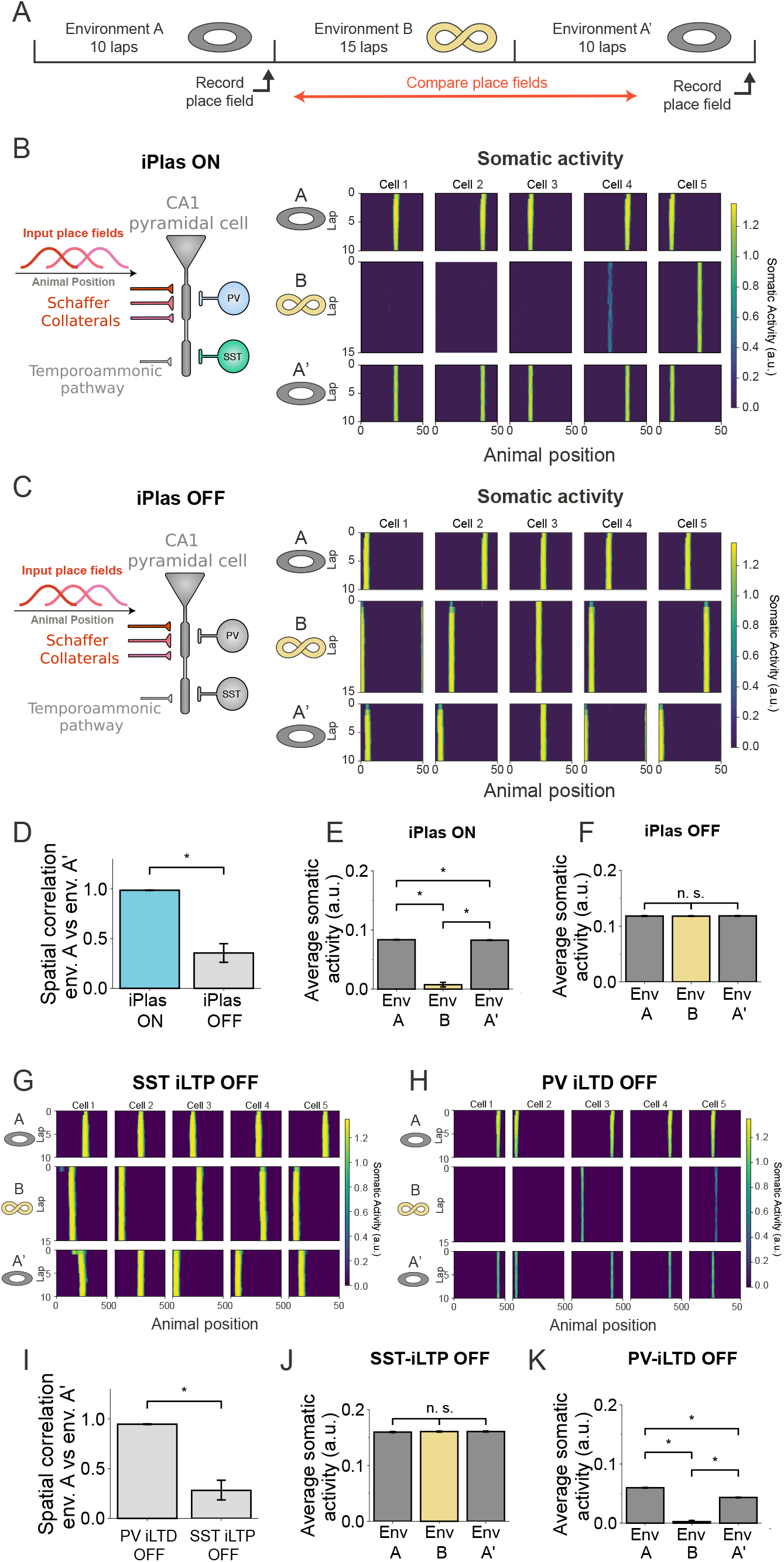
PV and SST plasticity ensure place cell stability and fidelity across multiple environments. **(A)** Simulation protocol. An animal explores environment A for 10 laps. It is then moved to environment B and explores it for 15 laps. Finally, the animal is moved back to the first environment (A’) and is allowed to run for another 10 laps. Throughout this protocol, two-compartment CA1 pyramidal cells are simulated receiving spatially-tuned Schaffer collateral inputs, temporoammonic input, and PV and SST inhibitory inputs. Schaffer collateral synapses follow a Hebbian-type excitatory plasticity dependent on the co-activation of dendritic and somatic compartments (see methods). PV and SST synapses undergo rate-based iPlas (PV-iLTD and SST-iLTP). We simulate the switch from environment A to environment B by randomly shuffling the identity of the SC inputs to the CA1 pyramidal neuron. Schematic depictions of environments A and B indicate their cyclical nature. **(B)** Diagram of simulated CA1 pyramidal cell and examples of somatic activity during exploration for iPlas ON. With iPlas (PV-iLTD and SST-iLTP) ON, place field location formed in environment A (top & bottom panel) remains stable after exposure to novel environment B (middle panel). **(C)** Diagram of simulated CA1 pyramidal cell and examples of somatic activity during exploration for iPlas OFF. Location of place fields formed in environment A are not maintained after exposure to a novel environment. **(D)** Spatial correlation between environment A before and after exposure to novel environment is maintained when iPlas is present but is reduced without iPlas. **(E)** When iPlas is ON, average somatic activity of recently formed place cell is significantly reduced in new environment B but returns to higher levels when the animal returns to environment A’. **(F)** When iPlas is OFF, average somatic activity remains high in both environments. **(G)** When SST-iLTP is turned off, leaving just PV-iLTD, place cell locations are not retained after exposure to a new environment. **(H)** When PV-iLTD is turned off, leaving just SST-iLTP, place cell locations are maintained across multiple environments. **(I)** Spatial correlation between environment A before and after exposure to novel environment is maintained when only PV-iLTD is turned off but is reduced when only SST-iLTP is turned off. **(J)** SST-iLTP is turned off, leaving just PV-iLTD, average somatic activity remains high in both environments. **(K)** When PV-iLTD is turned off, leaving just SST-iLTP, average somatic activity is lower and thus less robust. Data represent mean ± S.E.M statistical comparison via unpaired t-tests. Significant difference is indicated (*p < 0.05). See also Figure S6.

For any trial simulation of the network, the simulated CA1 pyramidal neuron rapidly developed a place field at a random location within environment A that remained stable for subsequent laps of the track (Figure 8B). These place fields were driven by rapidly evolving synaptic weight changes (Figure S6). When the track was switched to environment B, the indexes of Shaffer collateral inputs were shuffled. In this environment, the CA1 pyramidal neuron occasionally formed a new place field but was more often silent due to an inability to align and adapt synaptic weight increases after the inputs were shuffled (Figure 8B and Figure S6A). On returning to the familiar environment A’, the initial place field location was reinstated immediately (Figure 8B,D,E). These results are qualitatively in line with the experimentally observed physiology of place cell activity in different environments (Colgin et al., 2008).

To determine the role of inhibitory plasticity, we first removed PV-iLTD and SST-iLTP from the model. Simulations of this model lacking inhibitory plasticity showed similar place cell activity in environment A. In contrast, when the track was switched to environment B without inhibitory plasticity engaged, synaptic weight changes drove the generation of new place fields in every trial and the overall spiking rates were not reduced (Figure 8C and Figure S6B). Furthermore, on returning to environment A, the place fields were no longer reinstated but instead new representations evolved (Figure 8C,D,F). Thus, without inhibitory plasticity novel environments generate interference and the network is no longer capable of creating stable place field representations.

We next sought to distinguish the roles of PV-iLTD and SST-iLTP within this network. Simulations of a model with only PV-iLTD (SST-iLTP OFF) showed similar lack of place cell stability across environments A-B-A’ to simulations with no inhibitory plasticity and overall spiking rates were unchanged in environments B and A’ due to prior spiking rate saturation (Figure 8G,I,J). With implementation of only SST-iLTP (PV-iLTD OFF), place cell stability across environments A-B-A’ was reinstated but overall spiking rates were reduced compared to simulations with full inhibitory plasticity (Figure 8H,I,K).

These circuit-level modelling data show how long-term inhibitory plasticity can provide a mechanism for the experimentally observed phenomenon that newly formed place cells are stable with repeated exposure to an environment and don’t undergo interference from experiencing other environments. This stability is principally due to SST-iLTP which also reduces the efficiency of forming new place fields in different environments. The overall spike output of place cells is maintained by PV-iLTD which counteracts the reduction in spike output caused by SST-iLTP.

## Discussion

Inhibitory GABAergic synapses are known to undergo long-term plasticity but very few studies have defined which subpopulations of inhibitory interneurons are engaged and whether plasticity is induced by physiological firing patterns. This is important since distinct interneuron subtypes play highly specific roles within neuronal networks (Pelkey et al., 2017) and therefore plasticity at one inhibitory synapse may have very different effects to another. In this study we address this complexity and show that proximal and distal dendritically targeting interneuron synapses on CA1 pyramidal neurons have distinct plasticity rules within the hippocampus. These inhibitory synapses undergo homosynaptic plasticity in a Hebbian manner relying on the coincident activation of interneurons and pyramidal neurons (Figure 1,2). This coincident activity enables recruitment of VGCCs to provide local sources of Ca^2+^ able to alter inhibitory synapse strength (Figure 3,4,5).

By computationally modelling the effects of inhibitory plasticity at a single neuron level, we predicted that altered inhibition at distinct dendritic compartments dramatically alters pyramidal neuron output (Figure 6). We confirmed this experimentally showing action potential generation from proximally and dendritically targeting excitatory inputs is modulated by inhibitory plasticity in corresponding dendritic compartments (Figure 7).

By expanding our computational model, we show how plasticity at these distinct inhibitory synapses can play roles in the stabilisation of place cell activity within the hippocampus (Figure 8). This inhibitory plasticity stabilisation ensures place cell fidelity and resilience to interference from activity in multiple different environments.

### Inhibitory plasticity at distinct inhibitory synapses

The mechanisms and network implications of long-term plasticity at glutamatergic synapses have been extensively characterised. However, less attention has been paid to long-term plasticity at inhibitory synapses which are known to undergo dynamic changes in efficacy (Chiu et al., 2019; Kullmann et al., 2012; Pelkey et al., 2017). An array of unique mechanisms discovered for inhibitory plasticity suggests a lack of uniformity across the multiple subtype-specific inhibitory synapses.

Within the hippocampus, synapses from CCK expressing proximally targeting basket cells onto CA1 pyramidal neurons undergo an endocannabinoid mediated iLTD (Chevaleyre and Castillo, 2003). Here, mobilisation of retrograde endocannabinoid signalling results in long term suppression of GABA release (Chevaleyre and Castillo, 2003; Younts et al., 2016). We demonstrate that a similar morphological subtype of interneuron, the proximally targeting PV interneurons can also undergo iLTD but in an endocannabinoid independent mechanism (Figure 3), highlighting the diversity of plasticity mechanisms even among interneurons with similar morphology (Freund and Katona, 2007).

Other long-term inhibitory plasticity mechanisms include a persistent shift in the chloride reversal potential caused by coincident activity-dependent modulation of the KCC2 chloride transporter (Ormond and Woodin, 2011; Wang et al., 2006; Woodin et al., 2003). Interestingly, in the hippocampus this iLTD is reported in the feedforward inhibitory pathway for Schaffer collateral innervation of CA1, commensurate with PV basket cell innervation, and is dependent on L-type VGCC and NMDA receptor activation. An alternative set of mechanisms for PV synapse plasticity is reported in the auditory cortex where PV synapses undergo bidirectional iSTDP via BDNF and GABA_B_ dependent mechanisms (Vickers et al., 2018). However, these mechanisms do not appear to apply to PV-iLTD in the hippocampus and moreover, we found no evidence for PV-iLTP. The lack of PV-iLTP is also reported in the prefrontal cortex where SST but not PV synapses undergo iLTP (Chiu et al., 2018). This suggests that inhibitory plasticity rules may not be conserved across brain regions or that PV synapses undergo multiple forms of inhibitory plasticity. In contrast, the mechanism for iLTP at SST synapses may be broadly conserved, at least between prefrontal cortex and hippocampus, where activation of CAMKII via Ca^2+^ influx to dendrites is found to induce iLTP (Chiu et al., 2018). The mechanistic differences here relate to the source of Ca^2+^ influx, arising from NMDA receptors in prefrontal cortex and L- and T-type VGCCs in hippocampus. We further show that hippocampal SST synapses can be depressed by non-coincident pre- and post-synaptic spike timing and it will be interesting to find out if this is also the case for synapses in prefrontal cortex.

### Inhibitory plasticity relies on recruitment of T-type VGCC

We show that plasticity at both PV and SST synapses exhibits several key properties: (i) it depends on the coincident activity of inhibitory synapses and postsynaptic action potentials, (ii) it is synapse specific, and (iii) it relies on postsynaptic Ca^2+^ signalling. These properties are apparently contradictory since synapse-specific inhibition is hyperpolarising which usually inhibits Ca^2+^ influx and signalling. We show that this apparent contradiction is resolved by recruitment of T-type VGCCs. At resting membrane potentials, T-type VGCCs are in an inactivated state, which can be de-inactivated by a hyperpolarising membrane potential (Perez-Reyes, 2003; Randall and Tsien, 1997), this activation profile lends itself perfectly to recruitment by GABAergic synapse activity. Importantly, we show that pairing action potentials before inhibitory input or inhibitory input alone is insufficient to induce inhibitory plasticity. These observations suggest GABA synapse dependent de-inactivation of T-type VGCC is required prior to action potential activation of T-type VGCC, leading to a local synapse specific source of Ca^2+^ to drive inhibitory plasticity.

Indeed, there is considerable evidence linking GABA signalling and T-type VGCC activation. In the cerebellum and thalamus where T-type VGCCs are widely expressed, T-type VGCCs regulate inhibitory synapse strength (Aizenman et al., 1998; Pigeat et al., 2015; Sieber et al., 2013; Wang et al., 2006). In the thalamus GABAergic synapses onto thalamocortical neurons de-inactivate T-type VGCCs and reduce inhibitory synaptic strength (Pigeat et al., 2015). This thalamocortical inhibitory plasticity is also dependent on the interaction of calcineurin with GABA_A_ receptors, similar to findings in the hippocampus (Muir et al., 2010; Wang et al., 2003) and those we present here.

A striking finding in our results shows that identical induction protocols PV and SST synapses induce opposing forms of plasticity via the differential recruitment of calcineurin and CaMKII. Since CaMKII requires higher [Ca^2+^] to activate, the differential recruitment of calcineurin and CaMKII could be explained if postsynaptic [Ca^2+^] is higher at SST vs PV inhibitory synapses. In support of this hypothesis, we show that SST-iLTP relies on L-type as well as T-type VGCCs suggesting a higher level of Ca^2+^ entry. Alternatively, expression levels of VGCCs may increase at more distal dendritic locations and there is evidence that T-type VGCC expression is higher in dendritic regions of pyramidal neurons causing differential regulation of glutamatergic plasticity along the proximal-distal axis of pyramidal neurons (Isomura et al., 2002).

### The consequences of inhibitory synaptic plasticity on hippocampal network function

Our data support two separate functions of interneuron subtype-specific inhibitory plasticity on hippocampal network function. Firstly, increasing inhibitory inputs to distal dendritic locations on CA1 pyramidal neurons whilst reducing inhibition at proximal locations prioritises response to inputs from CA3 pyramidal neurons via the Schaffer collateral pathway over those from entorhinal neurons via the temporoammonic pathway. Secondly, in our computational model, increasing inhibition at distal dendritic locations inhibits the induction of synaptic plasticity at excitatory synapses (Leao et al., 2012; Williams and Holtmaat, 2019) thereby reducing adaptability of hippocampal representations.

Interneuron-specific inhibitory plasticity at proximal and distal dendritic locations coupled with the anatomical arrangement of Schaffer collateral inputs to proximal dendrites and temporoammonic inputs to distal dendrites intuitively predicts that inhibitory plasticity will rebalance the weighting of excitatory inputs in favour of Schaffer collaterals. We formalised these predictions using computational modelling of the CA1 network and then tested them experimentally. We confirmed that PV-iLTD increases CA1 pyramidal neuron responses to Schaffer collateral stimulation whereas SST-iLTP decreases responses to temporoammonic stimulation. Our combination of computational modelling and experimental approaches also showed that the majority of PV interneurons activated by our optogenetic approach are feedforward in the Schaffer collateral pathway but not the temporoammonic. Furthermore, our data indicate a limited feedback role for the PV interneurons we activate since PV-iLTD did not impact CA1 pyramidal neuron spike output in response to temporoammonic input. This broadly corresponds to anatomical and functional data for PV interneurons in the hippocampus (Booker and Vida, 2018; Ganter et al., 2004; Milstein et al., 2015; Sun et al., 2014). In contrast, the SST interneurons we stimulate are distally targeting, receive almost no feedforward input and are driven by feedback input from CA1 pyramidal neurons and therefore have all the hallmarks of OLM cells (Blasco-Ibanez and Freund, 1995; Booker and Vida, 2018; Lacaille et al., 1987; Sun et al., 2014).

The implications of reprioritising CA3 input over entorhinal input to CA1 are not straightforward but parallels can be drawn with the short-term reprioritisation caused by neuromodulator or thalamocortical inputs in cortical circuits (Fu et al., 2014; Hasselmo, 2006; Hasselmo and Schnell, 1994; Williams and Holtmaat, 2019). Often these mechanisms also involve reconfiguration of inhibitory interneuron circuits which are proposed to prioritise input of new sensory information over internal representation on short timescales including theta cycle timescales (Dupret et al., 2013; Lopes-Dos-Santos et al., 2018). The long-term inhibitory plasticity described here is predicted to achieve the reverse outcome prioritising previously learnt associations and making the network less receptive to new information.

Such a scenario would fit with the second role of inhibitory plasticity inhibiting excitatory plasticity and therefore the formation of new representations. SST inhibitory input regulates dendritic excitability and therefore NMDA receptor activation and excitatory synaptic plasticity (Schulz et al., 2018). Long-term plasticity of SST synapses will change the ability for CA1 pyramidal neurons to undergo induction of excitatory LTP (Leao et al., 2012; Williams and Holtmaat, 2019). We show that this has major implications for the stability and flexibility of place cells since their formation and remapping depends on excitatory synaptic plasticity driven by dendritic spikes generated by coincident Schaffer collateral and temporoammonic inputs (Bittner et al., 2015; Bittner et al., 2017; Sheffield and Dombeck, 2015). SST-iLTP prevents place cell representations in environment A being disrupted by different representations in environment B and indeed reduces the ability for place cells to be active in multiple environments. Interestingly, short-term changes in PV and SST interneuron firing rates in response to novel environments may provide a countermechanism to enable new place fields to be formed in novel environments (Sheffield et al., 2017). Our data and modelling therefore provide a mechanism to reconcile the observed stability of place cells across time and their ability to remap in distinct environments (Colgin et al., 2008).

In summary, our data reveal a novel form of inhibitory plasticity in the hippocampus induced by physiological patterns of firing. It has major implications for hippocampal function controlling input-output relationships in CA1 and providing a mechanism to explain a long-standing conundrum regarding place cell stability versus flexibility.

## Acknowledgements

We thank Rui Ponte Costa, David Dupret and Andrew Randall for critical input to previous versions of the manuscript and all members of the Clopath and Mellor groups for discussion. We also thank Klas Kullander for providing Chrna2-cre mice. This work was supported by Biotechnology and Biological Sciences Research Council (BBSRC), Wellcome Trust, the EPSRC and the Simons Foundation and NIH. V.P. is supported by CAPES Foundation, process n. 99999.001758/2015-02.

## Contributions

Conceptualization, C.C. and J.R.M.; Methodology, M.U., V.P. and S.E.L.C.; Investigation, M.U., V.P. and S.E.L.C.; Visualization, M.U. and V.P.; Writing – Original Draft, M.U., V.P., C.C. and J.R.M.; Writing – Review & Editing, M.U., V.P., C.C. and J.R.M.; Funding Acquisition, C.C. and J.R.M.; Supervision, C.C. and J.R.M.

## Methods

### Animal strains and breeding

All procedures and techniques were conducted in accordance to the UK animals scientific procedures act, 1986 with approval of the University of Bristol ethics committee. To express ChR2 within either PV,SST or Chrna2 expressing interneurons C57/Bl6 homozygous Ai32 mice (Gt(ROSA)26Sor^tm32(CAG-COP4*H134R/EYFP)Hze^ Jax Stock number: 024109) were bred with either homozygous PV-Cre (Pvalb^tm1(cre)Arbr/J^ Jax stock number: 017320), SST-Cre (Sst^tm2.1(cre)Zjh/J^ Jax stock number: 013044) or Chrna2-cre (Leao et al., 2012) mice creating heterozygous offspring with interneuron specific expression of ChR2. For brain slice electrophysiology both male and female mice were used.

### Brain slice preparation

Brain slices were prepared from 4-9 week old mice following cervical dislocation and decapitation and brains removed and dissected in ice cold cutting solution containing in mM: 205 Sucrose, 10 Glucose, 26 NaHCO_3_, 2.5 KCl, 1.25 NaH_2_PO_4_, 0.5 CaCl_2_, 5 MgCl_2_, constantly bubbled with 95% O_2_ and 5% CO_2_. Horizontal brain slices, 400 µM thick containing the hippocampus were prepared via a vibratome (Leica LS1200). Brain slices were transferred to ACSF containing in mM: 124 NaCl, 3 KCl, 24 NaHCO_3_, 1.25 NaH_2_PO_4_ 10 Glucose, 2.5 CaCl_2_, 1.3 MgCl_2_, constantly bubbled with 95% O_2_ and 5% CO_2._ and incubated at 35 °C for 30 min before being stored at room temperature for at least 30 min before experimentation.

### Whole cell patch clamp recordings

Brain slices were transferred to a submerged slice recording chamber with a constant 2.5 ml/min flow of ACSF (see above), held at 32 °C. Inhibitory plasticity experiments were recorded in the presence of DAP5 (50 µM) and NBQX (20 µM) to isolate GABAergic events. Slices were visualised using infrared DIC optics using a Scientifica SliceScope microscope. Patch electrodes with a resistance of 3-6 MΩ were pulled from borosilicate glass capillaries using a horizontal puller (P-97, Sutter-instruments) and filled with internal solution. For whole cell voltage clamp recordings where neurons were held at 0 mV internal solution consisted of in mM: 130 Cs-MeSO_4_, 4 NaCl, 10 HEPES, 0.5 EGTA, 10 TEA-Cl, 2 Mg-ATP, 0.5 Na2-GTP, 1 QX-314.Cl, adjusted to pH 7.3 with CsOH and ∼ 290 mOsm, Cl^-^ reversal potential −57 mV. For iSTDP experiments an intracellular solution consisting of in mM: 140 K-gluconate, 5 NaCl, 1 MgCl_2_, 10 HEPES, 4 Mg-ATP, 0.3 Na_2_-GTP, 0.2 EGTA adjusted to pH 7.3 with KOH and ∼ 290 mOsm, Cl^-^ reversal potential −77 mV. For current clamp recordings the intracellular solution consisted of in mM: 130 K-gluconate, 8 NaCl, 1 MgCl_2_, 10 HEPES, 4 Mg-ATP, 0.3 Na_2_-GTP, 0.2 EGTA adjusted to pH 7.3 with KOH and ∼290 mOsm, Cl^-^ reversal −67 mV. In all experiments a junction potential of ∼ −15 mV was not compensated.

Recordings of CA1 pyramidal neurons were conducted via a Multiclamp 700A amplifier (Molecular devices) filtered at 6 kHz and digitised at a sampling frequency of 20 kHz using a Micro 1401 data acquisition board (CED). Data was acquired using Signal5 software (CED) and data analysed using custom MATLAB Scripts.

### Synaptic stimulation and plasticity protocols

For inhibitory plasticity experiments subtype specific IPSCs were evoked via optical stimulation of ChR2 via a 470 nm LED (Thorlabs) through a 40x objective lens using 2-5 ms square pulses of light. Control pathway IPSCs were evoked via 100 µs square pulse electrical stimulation delivered via a monopolar stimulating electrode placed in the pyramidal layer or Stratum Radiatum. For plasticity experiments each pathway was stimulated every 15 sec in an interleaved fashion and synapses from each pathway were checked for independence by a paired pulse protocol (Figure S1).

For light evoked TBS plasticity, neurons were voltage clamped at 0 mV for the duration of the experiment, light evoked TBS was applied in voltage clamp and consisted of 5 bursts delivered at 5 hz, each burst containing 4 light pulses at 100 hz with the protocol repeated 5 times at 0.033 hz. For inhibitory spike time dependent plasticity experiments, neurons were voltage clamped at −50 mV. Pairing protocol consisted of 100 pairings at 5 hz in current clamp (neurons maintained at −50 mV), each consisting of presynaptic light stimulation with a burst of action potentials initiated via somatic current injections (2 ms duration, 1 nA amplitude). For all plasticity experiments the series resistance was monitored and cells that showed a >20% change were excluded from analysis.

Spike probability experiments were conducted in current clamp where 10 EPSPs were evoked at 10 hz via a bipolar stimulating electrode placed in either the SR or SLM layer to stimulate the Schaffer collateral or temporoammonic pathway. Stimulation intensity was adjusted to evoked action potentials in roughly half of the EPSP stimulations.

### Immunohistochemistry

Brains were fixed via cardiac perfusion of Phosphate buffered saline (PBS) followed by 4% Formaldehyde in PBS. Brains were removed and stored in PFA for 24hrs and then transferred to 30% sucrose PBS solution for 2 days. 50 µm thick slices were then obtained via cryostat sectioning. Slices were incubated in a blocking solution containing 5% donkey serum and 0.2% Triton X-100 for 90 min at room temperature. Slices were then incubated in room temperature overnight in PBS containing 1% donkey serum and either anti-PV (1:10000, Sigma P3088) anti-SST (1:10000 Santa Cruz SC-7819) or anti-GFP (1:1000 LifeTech A11122) antibodies for PV, SST and ChR2 visualisation respectively. Slices were then washed with PBS and incubated with secondary antibodies, Alexa-594 (1:1000, LifeTech) or Alexa-488 (1:1000, LifeTech) for 2hrs at room temperature, before washing with PBS and mounting on microscope slides with 1:1000 DAPI staining. Slices were then visualised, and images acquired using a widefield fluorescence microscope. Hippocampal layer regions within the CA1 were defined based on DAPI staining and ChR2 mean fluorescence intensity was quantified using ImageJ software.

### In vitro data analysis

Experimental unit was defined as cell with only one cell recorded per slice. Up to 3 cells were recorded from each animal with an average of 1.6 cells per animal. Measurements were taken as an average of 4 responses to obtain a data point per min, averages represent mean ± S.E.M. Time series data was normalised to the last 5 min of baseline and plasticity was assessed by comparing the average IPSC amplitude 20-30 min after plasticity induction between the control and test pathway. Owing to the within cell control, data were analysed using a paired two-tailed Student’s t-test between the two pathways. For histograms comparing the average test pathway plasticity, statistical significance was assigned as a one-sample t-test compared to 100. In all cases significance assigned if p < 0.05.

### Computational modelling

#### Neuron Model and network structure

We investigate a feedforward network consisted of a single postsynaptic neuron receiving inputs from the temporoammonic pathway, Schaffer collaterals, and SST and PV interneurons. The postsynaptic neuron is modelled using a two-compartment, rate-based neuron model. The first compartment represents the distal dendrites of CA1 pyramidal cells, receiving excitatory inputs from the temporoammonic pathway and inhibitory inputs from SST interneurons. The second compartment represents the perisomatic region of CA1 pyramidal cells, receiving excitatory inputs from Schaffer Collaterals and inhibitory inputs from PV interneurons.

The dendritic compartment’s activity, *r_dend_*, is given by

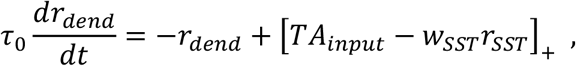

where [⋅]_+_ denotes a rectification that sets negative values to zero, *𝜏_0_* is a time constant, *TA_input_* is the temporoammonic pathway input, *w_SST_* is the synaptic weight from SST interneurons to the CA1 pyramidal cell, and *r_SST_* = 1 is the SST interneuron activity.

The somatic compartment’s activity, *r_soma_*, is given by

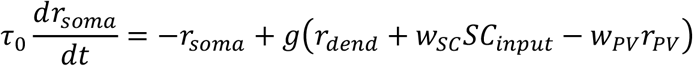

where *SC_input_* is the activity of Schaffer Collateral input neurons, *w_SC_* are the synaptic weights from SC inputs, *w_PV_* is the synaptic weight from PV interneurons to the CA1 pyramidal cell, *r_PV_* = 1 is the PV interneuron activity, and *g* is a non-linear function given by

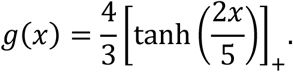

### TA and SC inputs

The simulated CA1 pyramidal cells receive excitatory inputs from the temporoammonic pathway and Schaffer Collaterals (SC). The input from the temporoammonic pathway is simulated as *TA_input_* = *μ_TA_* + *𝜉_TA_*, where *μ_TA_* is a constant and *𝜉_TA_* is generated from an Ornstein-Uhlenbeck process with a time constant of 50 ms, mean 0 and variance 0.5. The SC inputs are generated from *N_SC_* input neurons and each input neuron is tuned to a specific location such that their firing rates span over the entire environment. All place fields of SC input neurons have the same tuning width *𝜎_SC_* and amplitude *A_SC_*.

For the simulations involving exploration, the simulated animal explores an annular track of length *L* with speed *v*. The activity of a SC input neuron with place field centered at position *p*_0_ is

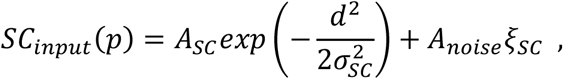

where *p* is the animal’s position, *d* is the distance between the *p* and *p*_0_ along the track, *A_noise_* is a constant, and *𝜉_SC_* is generated from an Ornstein-Uhlenbeck process with a time constant of 50 ms, mean 0 and variance 0.5.

In figure 6, we simulate an artificial stimulation of TA and SC. Therefore, for these simulations, SC inputs are not spatially tuned. Instead, they are simulated as *SC_input_* = *μ_SC_* + *𝜉_SC_*, where *μ_SC_* is a constant.

### Excitatory plasticity model

In figure 8, synaptic weights from SC input neurons to CA1 pyramidal neurons are plastic. These connections follow a Hebbian-type plasticity rule in which changes in synaptic weights depend on coincident pre- and postsynaptic activity. The postsynaptic term is given by the product of dendritic and somatic activity. This is motivated by recent findings suggesting that place fields are modified and formed following coincident TA and SC inputs (Bittner et al., 2015; Bittner et al., 2017). The excitatory synaptic weight *w_ij_* from input neuron *j* to postsynaptic neuron *i* is updated following

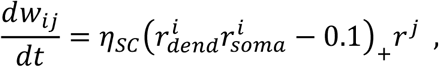

where 𝜂*_SC_* is the learning rate for SC connections, *r^j^* is the presynaptic neuron activity, 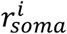 is the somatic activity of the postsynaptic neuron, and 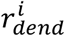 is the dendritic activity of the postsynaptic neuron. Because this rule is inherently unstable, synaptic weights are also normalized as commonly done (Lazar et al., 2009). After every weight update, we subtract the average synaptic weight ∑*_j_ w_ij_* /*N_SC_* from all weights and add a constant term (here 2). Negative weights are then rectified to zero.

### Inhibitory plasticity model

We implement an inhibitory plasticity rule inspired by our experimental findings. Under a rate-based framework, these plasticity rules are assumed to mirror those found during theta oscillations. Synaptic weights from PV interneurons onto CA1 pyramidal cells follow a rate-based Hebbian plasticity rule in which the coactivation of pre- and postsynaptic neurons leads to LTD:

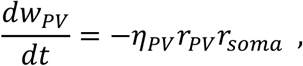

where 𝜂*_PV_* is the learning rate for PV connections, and *r_soma_* is pyramidal cell somatic activity. Synaptic weights from SST interneurons onto CA1 pyramidal cells follow a Hebbian plasticity rule in which the coactivation of pre- and postsynaptic neurons leads to LTP:

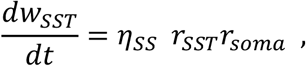

where 𝜂*_SST_* is the learning rate for SST connections. Both PV and SST synaptic weights are bounded between *w_min_* and *w_max_*.

### Environment switch

In figure 8, we simulate a feedforward network while an animal runs through an annular track. When the animal starts exploring environment A for the first time, the initial synaptic weights for the SC inputs are drawn from a lognormal distribution with underlying normal distribution with mean zero and standard deviation 0.1. The synaptic weights are then multiplied by 0.1 and two neighbouring inputs are randomly chosen and their synaptic weights are set to 0.6. This imposes a small structure in the synaptic weights and ensures that input neurons are able to induce postsynaptic activity. The animal then explores environment A for 10 laps. Next, the animal is moved to a novel environment (environment B). We simulate the switch to a novel environment by randomly shuffling the identity of the SC inputs to the CA1 pyramidal neuron. The animal then explores environment B for 15 laps. Subsequently, the animal is moved back to environment A (environment A’), which is implemented by returning the SC inputs to the original identity. Finally, the animal explores environment A’ for another 10 laps.

### Parameters and simulations

All simulations were implemented in python and are available at ModelDB. The parameters used in our simulations can be found in table 1.

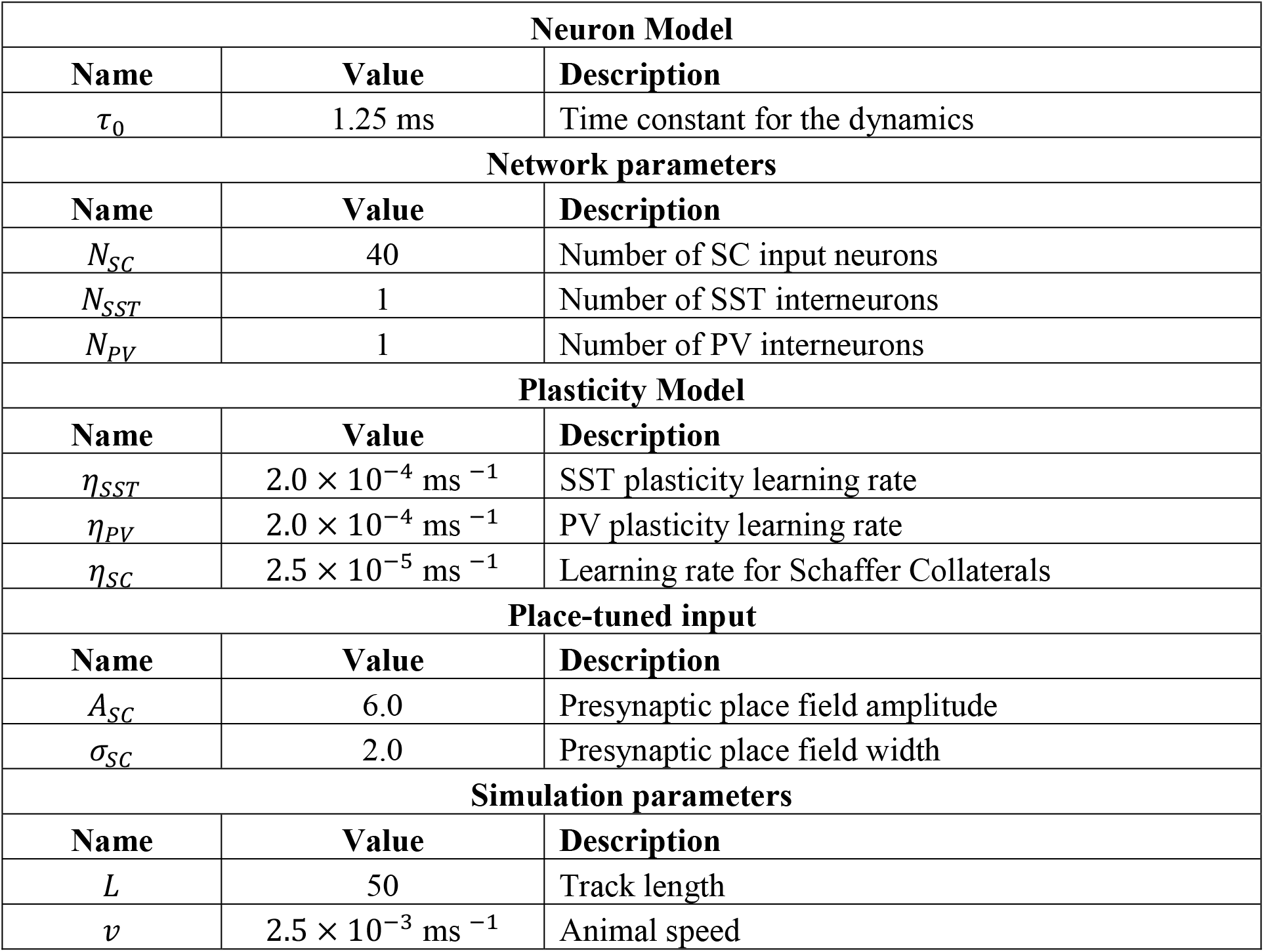

---

**Figure S1 (related to Figure 1).**
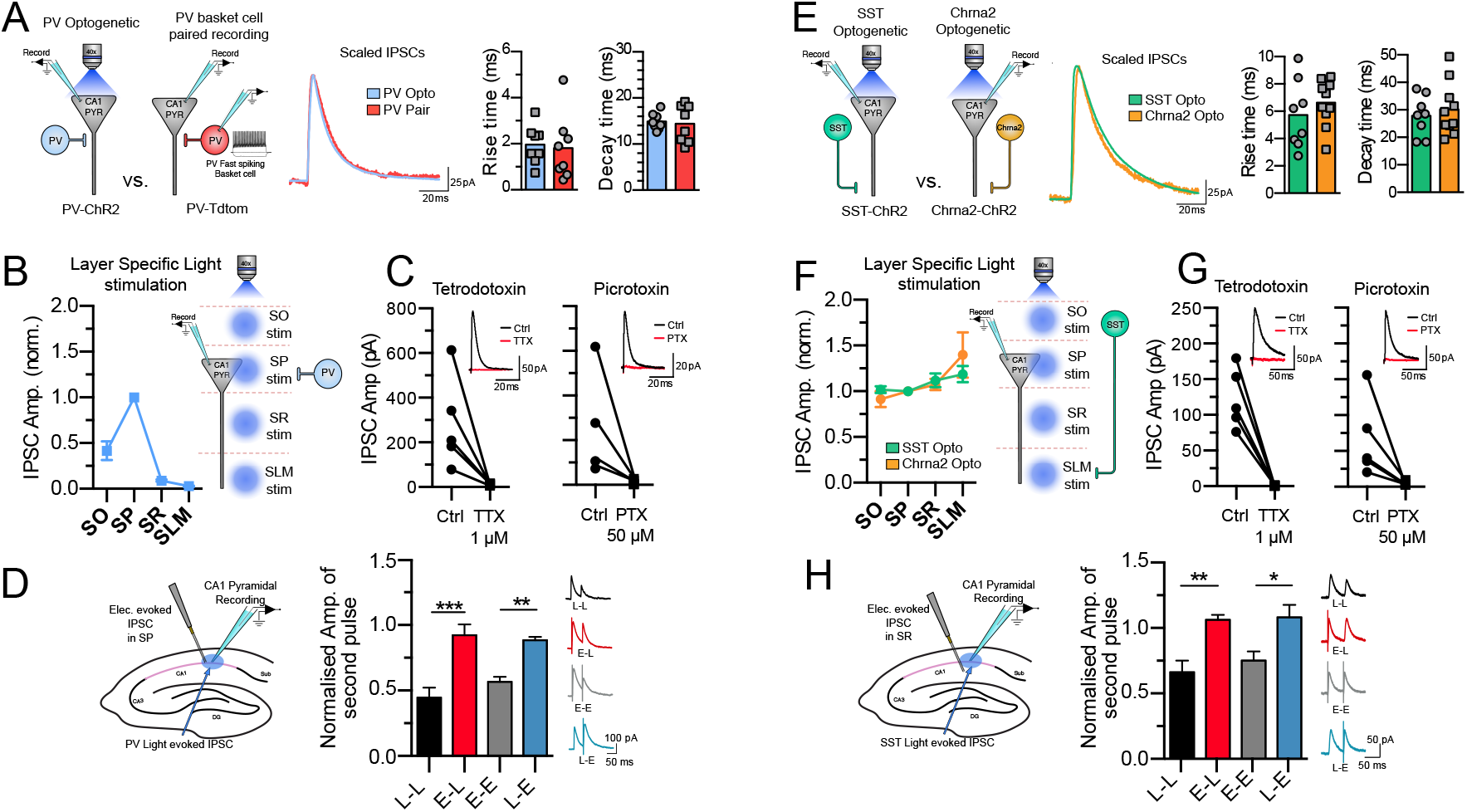
Characterisation of PV and SST optogenetic responses. **(A)** Comparison of optogenetically evoked PV IPSCs and IPSCs recorded via PV fast-spiking basket cell and CA1 pyramidal neuron paired recordings. Both rise and decay times were indistinguishable. **(B)** PV IPSCs evoked via light stimulation through the objective lens positioned over different layers of the hippocampus. PV IPSC amplitude was highest in the pyramidal layer consistent with PV basket cell activation. **(C)** PV IPSCs were completely blocked by 1µM tetrodotoxin and 50µM picrotoxin showing optogenetic IPSCs are action potential and GABAA receptor dependent. **(D)** Check of pathway independence between PV-optogenetic light-evoked pathway and electrically-evoked IPSC control pathway in SP. Electrical stimulation failed to depress light responses whilst light responses failed to depress electrical responses indicating separate discrete inhibitory synapse activation. **(E)** Comparison of optogenetically evoked IPSCs SST and Chrna2 expressing OLM interneurons. Both rise and decay times were indistinguishable. **(F)** SST and Chrna2 IPSCs evoked via light stimulation of different layers of the hippocampus. Both SST and Chrna2 IPSC amplitudes were maintained across all layers and highest in stratum lacunosum moleculare layer consistent with SST OLM interneuron activation. **(G)** SST IPSCs were completely blocked by 1µM tetrodotoxin and 50µM picrotoxin showing optogenetic IPSCs are action potential and GABAA receptor dependent. **(H)** Check of pathway independence between SST-optogenetic light-evoked pathway and electrically-evoked IPSC control pathway in the SR. Electrical stimulation failed to depress light responses whilst light responses failed to depress electrical responses indicating separate discrete inhibitory synapse activation. Data represent mean ± S.E.M statistical comparison via unpaired t-tests (D,H) (*p < 0.05, **p < 0.01, ***p < 0.001).

**Figure S2 (related to Figure 1).**
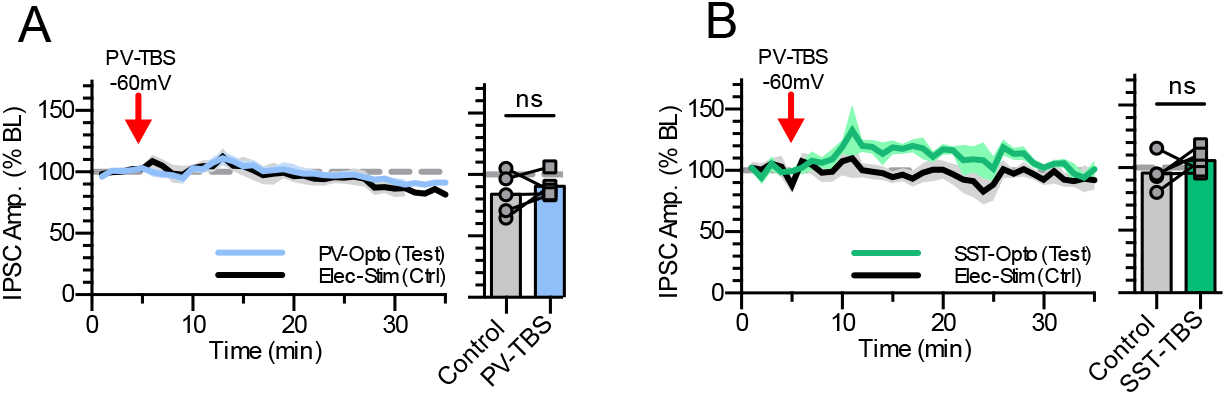
TBS induced PV-iLTD and SST-iLTP are dependent on postsynaptic depolarisation. CA1 pyramidal neurons were recorded at 0 mV for the duration of the experiment except for during the light induced TBS protocol in which the neuron was held at −60 mV **(A)** PV-iLTD was not induced if CA1 pyramidal neurons were held at −60 mV during the induction protocol. **(B)** SST-iLTP failed to be induced when CA1 pyramidal neurons were held at −60 mV during the induction protocol. Data represent mean ± S.E.M statistical comparison via paired t-tests.

**Figure S3 (related to Figure 3).**
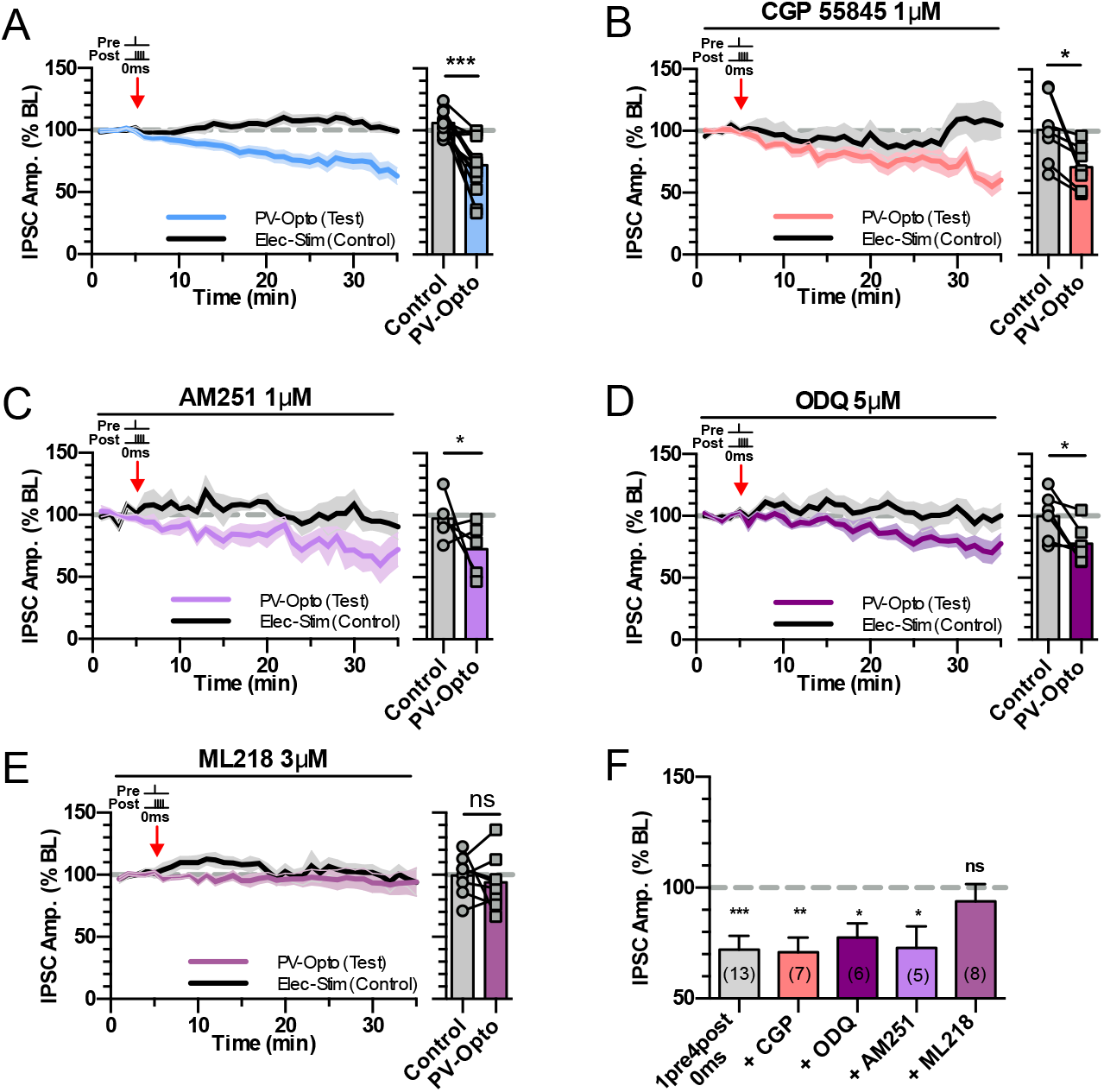
PV-iLTD is not dependent on endocannabinoid, GABAB receptors or nitrous oxide signalling. **(A)** PV-iLTD induced by 0 ms 1pre 4post timing (data from Figure 2C). **(B)** GABAB antagonist CGP55845 (1 µM) failed to block PV-iLTD **(C)** CB1 receptor antagonist AM251 (1 µM) failed to block PV-iLTD. **(D)** Inhibiting the Nitrous oxide pathway via inhibition of guanylyl cyclase with ODQ (5 µM) failed to block PV-iLTD. **(E)** The selective T-type VGCC inhibitor ML218 (3 µM) blocked PV-iLTD **(F)** Summary histogram displaying the level of plasticity under each experimental condition. Data represent mean ± S.E.M statistical comparison via paired t-tests (A-E) and one sample t-tests (F). Significant difference is indicated (*p < 0.05).

**Figure S4 (related to Figure 4).**
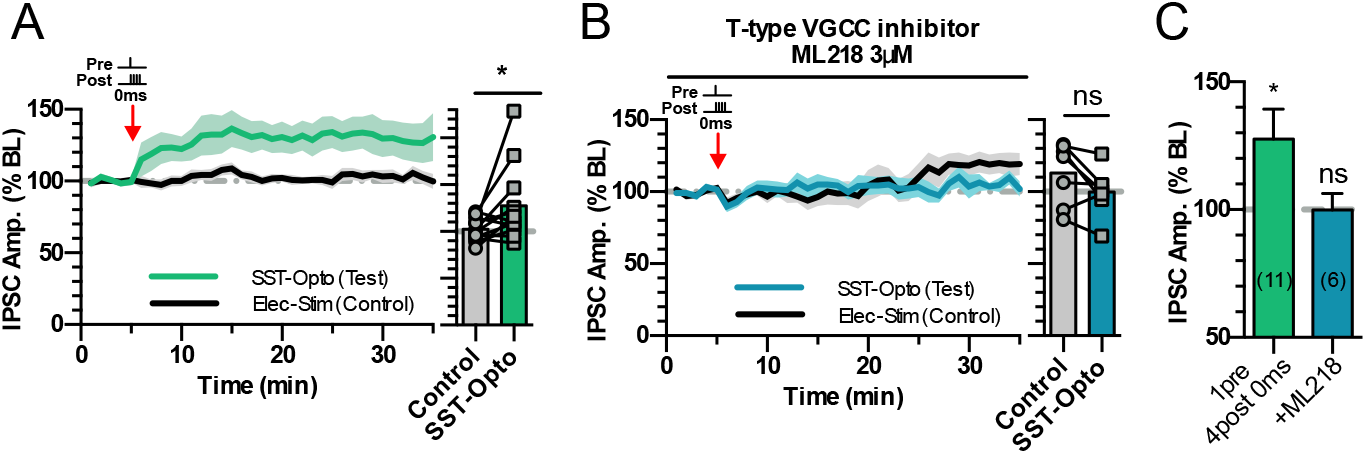
SST-iLTP requires T-type VGCC activation. **(A)** SST-iLTP induced by 0 ms 1pre 4post timing (data from Figure 2G). **(B)** The selective T-type VGCC inhibitor ML218 (3 µM) blocked SST-iLTP **(C)** Summary histogram displaying the level of plasticity under each experimental condition. Data represent mean ± S.E.M statistical comparison via paired t-tests (A,B) and one sample t-tests (C). Significant difference is indicated (*p < 0.05).

**Figure S5 (related to Figure 6).**
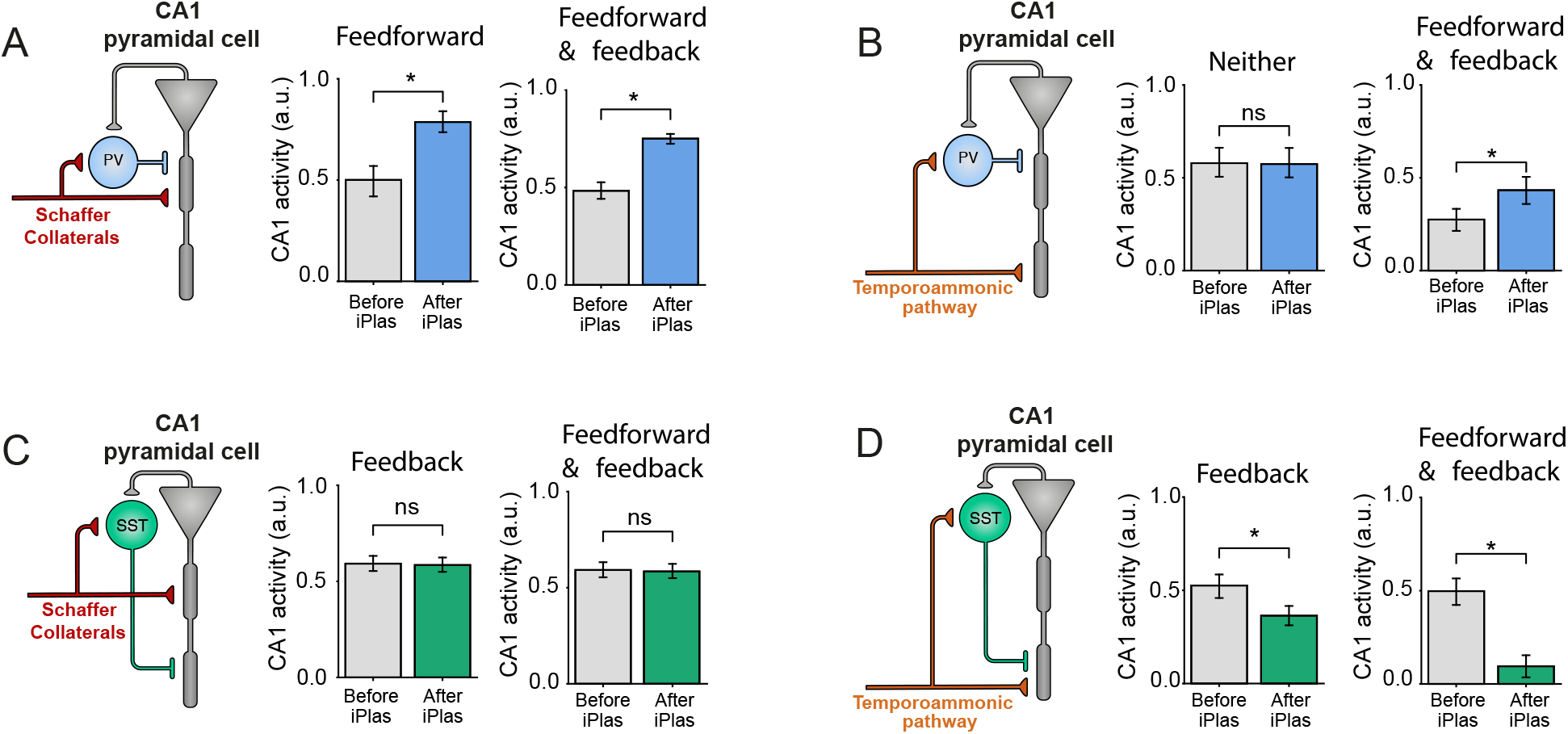
PV and SST plasticity regulate CA1 pyramidal neuron excitability via feedback and feedforward inhibition. **(A)** PV-iLTD increased CA1 pyramidal neuron activity in responses to Schaffer collateral input if PV interneurons are engaged via feedforward inhibition or feedforward and feedback inhibition. **(B)** PV-iLTD had no effect on temporoammonic excitation of CA1 pyramidal neurons due to lack of feedforward or feedback inhibition. If the temporoammonic pathway recruits PV interneurons via feedforward or partake in feedback inhibition PV-iLTD increased temporoammonic pathway driven CA1 activity. **(C)** SST-iLTP had no effect on the Schaffer collateral induced CA1 pyramidal neuron excitability if it is engaged via feedback or feedback and feedforward inhibition. **(D)** SST-iLTP reduced the temporoammonic pathway driven excitability of CA1 pyramidal neurons when engaged via feedback or feedforward and feedback inhibition. Data represent mean ± S.E.M statistical comparison via unpaired t-tests. Significant difference is indicated (*p < 0.05).

**Figure S6 (related to Figure 8).**
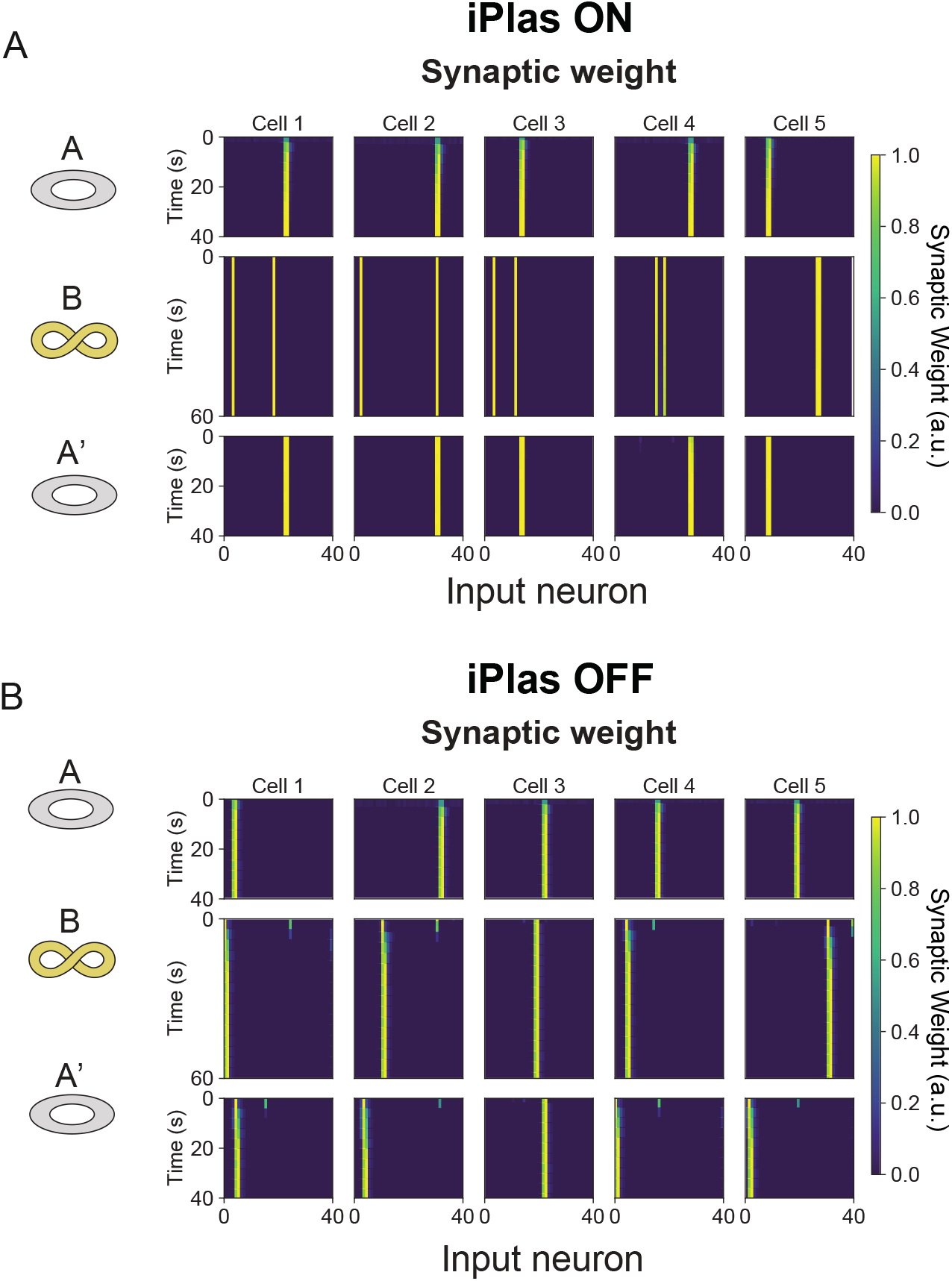
Simulated synaptic weight evolution during exploration in different environments. **(A)** Evolution of synaptic weights over time for the example cells shown in Figure 8B. During these simulations, iPlas is active. **(B)** Evolution of synaptic weights over time for the example cells shown in Figure 8C. During these simulations, iPlas is turned off.

## References

Aizenman, C.D., Manis, P.B., and Linden, D.J. (1998). Polarity of long-term synaptic gain change is related to postsynaptic spike firing at a cerebellar inhibitory synapse. Neuron 21, 827–835.

Alger, B.E., and Pitler, T.A. (1995). Retrograde signaling at GABAA-receptor synapses in the mammalian CNS. Trends Neurosci 18, 333–340.

Bittner, K.C., Grienberger, C., Vaidya, S.P., Milstein, A.D., Macklin, J.J., Suh, J., Tonegawa, S., and Magee, J.C. (2015). Conjunctive input processing drives feature selectivity in hippocampal CA1 neurons. Nat Neurosci 18, 1133–1142.

Bittner, K.C., Milstein, A.D., Grienberger, C., Romani, S., and Magee, J.C. (2017). Behavioral time scale synaptic plasticity underlies CA1 place fields. Science 357, 1033–1036.

Blasco-Ibanez, J.M., and Freund, T.F. (1995). Synaptic input of horizontal interneurons in stratum oriens of the hippocampal CA1 subfield: structural basis of feed-back activation. Eur J Neurosci 7, 2170–2180.

Booker, S.A., and Vida, I. (2018). Morphological diversity and connectivity of hippocampal interneurons. Cell Tissue Res 373, 619–641.

Buzsaki, G. (2002). Theta oscillations in the hippocampus. Neuron 33, 325–340.

Chaudhuri, R., and Fiete, I. (2016). Computational principles of memory. Nat Neurosci 19, 394–403.

Chevaleyre, V., and Castillo, P.E. (2003). Heterosynaptic LTD of hippocampal GABAergic synapses: a novel role of endocannabinoids in regulating excitability. Neuron 38, 461–472.

Chiu, C.Q., Barberis, A., and Higley, M.J. (2019). Preserving the balance: diverse forms of long-term GABAergic synaptic plasticity. Nat Rev Neurosci 20, 272–281.

Chiu, C.Q., Lur, G., Morse, T.M., Carnevale, N.T., Ellis-Davies, G.C., and Higley, M.J. (2013). Compartmentalization of GABAergic inhibition by dendritic spines. Science 340, 759–762.

Chiu, C.Q., Martenson, J.S., Yamazaki, M., Natsume, R., Sakimura, K., Tomita, S., Tavalin, S.J., and Higley, M.J. (2018). Input-Specific NMDAR-Dependent Potentiation of Dendritic GABAergic Inhibition. Neuron 97, 368–377 e363.

Cohen, J.D., Bolstad, M., and Lee, A.K. (2017). Experience-dependent shaping of hippocampal CA1 intracellular activity in novel and familiar environments. Elife 6.

Colgin, L.L., Moser, E.I., and Moser, M.B. (2008). Understanding memory through hippocampal remapping. Trends Neurosci 31, 469–477.

Del Pino, I., Brotons-Mas, J.R., Marques-Smith, A., Marighetto, A., Frick, A., Marin, O., and Rico, B. (2017). Abnormal wiring of CCK(+) basket cells disrupts spatial information coding. Nat Neurosci 20, 784–792.

Donato, F., Rompani, S.B., and Caroni, P. (2013). Parvalbumin-expressing basket-cell network plasticity induced by experience regulates adult learning. Nature 504, 272–276.

Dupret, D., O’Neill, J., and Csicsvari, J. (2013). Dynamic reconfiguration of hippocampal interneuron circuits during spatial learning. Neuron 78, 166–180.

Freund, T.F., and Katona, I. (2007). Perisomatic inhibition. Neuron 56, 33–42.

Fu, Y., Tucciarone, J.M., Espinosa, J.S., Sheng, N., Darcy, D.P., Nicoll, R.A., Huang, Z.J., and Stryker, M.P. (2014). A cortical circuit for gain control by behavioral state. Cell 156, 1139–1152.

Ganter, P., Szucs, P., Paulsen, O., and Somogyi, P. (2004). Properties of horizontal axo-axonic cells in stratum oriens of the hippocampal CA1 area of rats in vitro. Hippocampus 14, 232–243.

Geiger, J.R., Lubke, J., Roth, A., Frotscher, M., and Jonas, P. (1997). Submillisecond AMPA receptor-mediated signaling at a principal neuron-interneuron synapse. Neuron 18, 1009–1023.

Griffith, T., Tsaneva-Atanasova, K., and Mellor, J.R. (2016). Control of Ca2+ Influx and Calmodulin Activation by SK-Channels in Dendritic Spines. PLoS computational biology 12, e1004949.

Harris, K.D., Hochgerner, H., Skene, N.G., Magno, L., Katona, L., Bengtsson Gonzales, C., Somogyi, P., Kessaris, N., Linnarsson, S., and Hjerling-Leffler, J. (2018). Classes and continua of hippocampal CA1 inhibitory neurons revealed by single-cell transcriptomics. PLoS biology 16, e2006387.

Hasselmo, M.E. (2006). The role of acetylcholine in learning and memory. Curr Opin Neurobiol 16, 710–715.

Hasselmo, M.E., and Schnell, E. (1994). Laminar selectivity of the cholinergic suppression of synaptic transmission in rat hippocampal region CA1: computational modeling and brain slice physiology. J Neurosci 14, 3898–3914.

Hellyer, P.J., Jachs, B., Clopath, C., and Leech, R. (2016). Local inhibitory plasticity tunes macroscopic brain dynamics and allows the emergence of functional brain networks. NeuroImage 124, 85–95.

Horn, M.E., and Nicoll, R.A. (2018). Somatostatin and parvalbumin inhibitory synapses onto hippocampal pyramidal neurons are regulated by distinct mechanisms. Proc Natl Acad Sci U S A 115, 589–594.

Isaac, J.T., Buchanan, K.A., Muller, R.U., and Mellor, J.R. (2009). Hippocampal place cell firing patterns can induce long-term synaptic plasticity in vitro. J Neurosci 29, 6840–6850.

Isomura, Y., Fujiwara-Tsukamoto, Y., Imanishi, M., Nambu, A., and Takada, M. (2002). Distance-dependent Ni(2+)-sensitivity of synaptic plasticity in apical dendrites of hippocampal CA1 pyramidal cells. J Neurophysiol 87, 1169–1174.

Katona, L., Lapray, D., Viney, T.J., Oulhaj, A., Borhegyi, Z., Micklem, B.R., Klausberger, T., and Somogyi, P. (2014). Sleep and movement differentiates actions of two types of somatostatin-expressing GABAergic interneuron in rat hippocampus. Neuron 82, 872–886.

Klausberger, T., Magill, P.J., Marton, L.F., Roberts, J.D., Cobden, P.M., Buzsaki, G., and Somogyi, P. (2003). Brain-state- and cell-type-specific firing of hippocampal interneurons in vivo. Nature 421, 844–848.

Klausberger, T., and Somogyi, P. (2008). Neuronal diversity and temporal dynamics: the unity of hippocampal circuit operations. Science 321, 53–57.

Kullmann, D.M., Moreau, A.W., Bakiri, Y., and Nicholson, E. (2012). Plasticity of inhibition. Neuron 75, 951–962.

Lacaille, J.C., Mueller, A.L., Kunkel, D.D., and Schwartzkroin, P.A. (1987). Local circuit interactions between oriens/alveus interneurons and CA1 pyramidal cells in hippocampal slices: electrophysiology and morphology. J Neurosci 7, 1979–1993.

Lazar, A., Pipa, G., and Triesch, J. (2009). SORN: a self-organizing recurrent neural network. Frontiers in computational neuroscience 3.

Leao, R.N., Mikulovic, S., Leao, K.E., Munguba, H., Gezelius, H., Enjin, A., Patra, K., Eriksson, A., Loew, L.M., Tort, A.B., and Kullander, K. (2012). OLM interneurons differentially modulate CA3 and entorhinal inputs to hippocampal CA1 neurons. Nat Neurosci 15, 1524–1530.

Lee, S.H., Foldy, C., and Soltesz, I. (2010). Distinct endocannabinoid control of GABA release at perisomatic and dendritic synapses in the hippocampus. J Neurosci 30, 7993–8000.

Lopes-Dos-Santos, V., van de Ven, G.M., Morley, A., Trouche, S., Campo-Urriza, N., and Dupret, D. (2018). Parsing Hippocampal Theta Oscillations by Nested Spectral Components during Spatial Exploration and Memory-Guided Behavior. Neuron 100, 940–952 e947.

Luscher, B., Fuchs, T., and Kilpatrick, C.L. (2011). GABAA receptor trafficking-mediated plasticity of inhibitory synapses. Neuron 70, 385–409.

Maccaferri, G. (2005). Stratum oriens horizontal interneurone diversity and hippocampal network dynamics. J Physiol 562, 73–80.

Magee, J.C., and Johnston, D. (1997). A synaptically controlled, associative signal for Hebbian plasticity in hippocampal neurons. Science 275, 209–213.

Marsden, K.C., Beattie, J.B., Friedenthal, J., and Carroll, R.C. (2007). NMDA receptor activation potentiates inhibitory transmission through GABA receptor-associated protein-dependent exocytosis of GABA(A) receptors. J Neurosci 27, 14326–14337.

Mikulovic, S., Restrepo, C.E., Hilscher, M.M., Kullander, K., and Leao, R.N. (2015). Novel markers for OLM interneurons in the hippocampus. Front Cell Neurosci 9, 201.

Milstein, A.D., Bloss, E.B., Apostolides, P.F., Vaidya, S.P., Dilly, G.A., Zemelman, B.V., and Magee, J.C. (2015). Inhibitory Gating of Input Comparison in the CA1 Microcircuit. Neuron 87, 1274–1289.

Mishra, R.K., Kim, S., Guzman, S.J., and Jonas, P. (2016). Symmetric spike timing-dependent plasticity at CA3-CA3 synapses optimizes storage and recall in autoassociative networks. Nature communications 7, 11552.

Muir, J., Arancibia-Carcamo, I.L., MacAskill, A.F., Smith, K.R., Griffin, L.D., and Kittler, J.T. (2010). NMDA receptors regulate GABAA receptor lateral mobility and clustering at inhibitory synapses through serine 327 on the gamma2 subunit. Proc Natl Acad Sci U S A 107, 16679–16684.

Nugent, F.S., Penick, E.C., and Kauer, J.A. (2007). Opioids block long-term potentiation of inhibitory synapses. Nature 446, 1086–1090.

Nusser, Z., Hajos, N., Somogyi, P., and Mody, I. (1998). Increased number of synaptic GABA(A) receptors underlies potentiation at hippocampal inhibitory synapses. Nature 395, 172–177.

O’Keefe, J. (1976). Place units in the hippocampus of the freely moving rat. Exp Neurol 51, 78–109.

Ormond, J., and Woodin, M.A. (2011). Disinhibition-Mediated LTP in the Hippocampus is Synapse Specific. Front Cell Neurosci 5, 17.

Pelkey, K.A., Chittajallu, R., Craig, M.T., Tricoire, L., Wester, J.C., and McBain, C.J. (2017). Hippocampal GABAergic Inhibitory Interneurons. Physiol Rev 97, 1619–1747.

Perez-Reyes, E. (2003). Molecular physiology of low-voltage-activated t-type calcium channels. Physiol Rev 83, 117–161.

Petrini, E.M., Ravasenga, T., Hausrat, T.J., Iurilli, G., Olcese, U., Racine, V., Sibarita, J.B., Jacob, T.C., Moss, S.J., Benfenati, F., et al. (2014). Synaptic recruitment of gephyrin regulates surface GABAA receptor dynamics for the expression of inhibitory LTP. Nature communications 5, 3921.

Pigeat, R., Chausson, P., Dreyfus, F.M., Leresche, N., and Lambert, R.C. (2015). Sleep slow wave-related homo and heterosynaptic LTD of intrathalamic GABAAergic synapses: involvement of T-type Ca2+ channels and metabotropic glutamate receptors. J Neurosci 35, 64–73.

Pouille, F., and Scanziani, M. (2001). Enforcement of temporal fidelity in pyramidal cells by somatic feed-forward inhibition. Science 293, 1159–1163.

Randall, A.D., and Tsien, R.W. (1997). Contrasting biophysical and pharmacological properties of T-type and R-type calcium channels. Neuropharmacology 36, 879–893.

Royer, S., Zemelman, B.V., Losonczy, A., Kim, J., Chance, F., Magee, J.C., and Buzsaki, G. (2012). Control of timing, rate and bursts of hippocampal place cells by dendritic and somatic inhibition. Nat Neurosci 15, 769–775.

Sadowski, J.H., Jones, M.W., and Mellor, J.R. (2016). Sharp-Wave Ripples Orchestrate the Induction of Synaptic Plasticity during Reactivation of Place Cell Firing Patterns in the Hippocampus. Cell reports 14, 1916–1929.

Schuemann, A., Klawiter, A., Bonhoeffer, T., and Wierenga, C.J. (2013). Structural plasticity of GABAergic axons is regulated by network activity and GABAA receptor activation. Front Neural Circuits 7, 113.

Schulz, J.M., Knoflach, F., Hernandez, M.C., and Bischofberger, J. (2018). Dendrite-targeting interneurons control synaptic NMDA-receptor activation via nonlinear alpha5-GABAA receptors. Nature communications 9, 3576.

Sheffield, M.E., and Dombeck, D.A. (2015). Calcium transient prevalence across the dendritic arbour predicts place field properties. Nature 517, 200–204.

Sheffield, M.E.J., Adoff, M.D., and Dombeck, D.A. (2017). Increased Prevalence of Calcium Transients across the Dendritic Arbor during Place Field Formation. Neuron 96, 490–504 e495.

Sieber, A.R., Min, R., and Nevian, T. (2013). Non-Hebbian long-term potentiation of inhibitory synapses in the thalamus. J Neurosci 33, 15675–15685.

Sohal, V.S., Zhang, F., Yizhar, O., and Deisseroth, K. (2009). Parvalbumin neurons and gamma rhythms enhance cortical circuit performance. Nature 459, 698–702.

Sun, Y., Nguyen, A.Q., Nguyen, J.P., Le, L., Saur, D., Choi, J., Callaway, E.M., and Xu, X. (2014). Cell-type-specific circuit connectivity of hippocampal CA1 revealed through Cre-dependent rabies tracing. Cell reports 7, 269–280.

Tigaret, C.M., Olivo, V., Sadowski, J.H., Ashby, M.C., and Mellor, J.R. (2016). Coordinated activation of distinct Ca(2+) sources and metabotropic glutamate receptors encodes Hebbian synaptic plasticity. Nature communications 7, 10289.

Trouche, S., Perestenko, P.V., van de Ven, G.M., Bratley, C.T., McNamara, C.G., Campo-Urriza, N., Black, S.L., Reijmers, L.G., and Dupret, D. (2016). Recoding a cocaine-place memory engram to a neutral engram in the hippocampus. Nat Neurosci 19, 564–567.

Varga, C., Golshani, P., and Soltesz, I. (2012). Frequency-invariant temporal ordering of interneuronal discharges during hippocampal oscillations in awake mice. Proc Natl Acad Sci U S A 109, E2726–2734.

Vickers, E.D., Clark, C., Osypenko, D., Fratzl, A., Kochubey, O., Bettler, B., and Schneggenburger, R. (2018). Parvalbumin-Interneuron Output Synapses Show Spike-Timing-Dependent Plasticity that Contributes to Auditory Map Remodeling. Neuron 99, 720–735 e726.

Vogels, T.P., Sprekeler, H., Zenke, F., Clopath, C., and Gerstner, W. (2011). Inhibitory plasticity balances excitation and inhibition in sensory pathways and memory networks. Science 334, 1569–1573.

Wang, J., Liu, S., Haditsch, U., Tu, W., Cochrane, K., Ahmadian, G., Tran, L., Paw, J., Wang, Y., Mansuy, I., et al. (2003). Interaction of calcineurin and type-A GABA receptor gamma 2 subunits produces long-term depression at CA1 inhibitory synapses. J Neurosci 23, 826–836.

Wang, L., Kitai, S.T., and Xiang, Z. (2006). Activity-dependent bidirectional modification of inhibitory synaptic transmission in rat subthalamic neurons. J Neurosci 26, 7321–7327.

Williams, L.E., and Holtmaat, A. (2019). Higher-Order Thalamocortical Inputs Gate Synaptic Long-Term Potentiation via Disinhibition. Neuron 101, 91–102 e104.

Woodin, M.A., Ganguly, K., and Poo, M.M. (2003). Coincident pre- and postsynaptic activity modifies GABAergic synapses by postsynaptic changes in Cl-transporter activity. Neuron 39, 807–820.

Xiang, Z., Thompson, A.D., Brogan, J.T., Schulte, M.L., Melancon, B.J., Mi, D., Lewis, L.M., Zou, B., Yang, L., Morrison, R., et al. (2011). The Discovery and Characterization of ML218: A Novel, Centrally Active T-Type Calcium Channel Inhibitor with Robust Effects in STN Neurons and in a Rodent Model of Parkinson’s Disease. ACS Chem Neurosci 2, 730–742.

Younts, T.J., Monday, H.R., Dudok, B., Klein, M.E., Jordan, B.A., Katona, I., and Castillo, P.E. (2016). Presynaptic Protein Synthesis Is Required for Long-Term Plasticity of GABA Release. Neuron 92, 479–492.

Ziv, Y., Burns, L.D., Cocker, E.D., Hamel, E.O., Ghosh, K.K., Kitch, L.J., El Gamal, A., and Schnitzer, M.J. (2013). Long-term dynamics of CA1 hippocampal place codes. Nat Neurosci 16, 264–266.

